# Allometric derivation and estimation of *Guadua weberbaueri* and *G. sarcocarpa* biomass in the bamboo-dominated forests of SW Amazonia

**DOI:** 10.1101/129262

**Authors:** Noah Yavit

## Abstract

Bamboo-dominated forests in Southwestern Amazonia encompass an estimated 180,000 km^2^ of nearly contiguous primary, tropical lowland forest. This area, largely composed of two bamboo species, *Guadua weberbaueri* Pilger and *G. sarcocarpa* Londoño & Peterson, comprises a significant portion of the Amazon Basin and has a potentially important effect on regional carbon storage. Numerous local REDD(+) projects would benefit from the development of allometric models for these species, although there has been just one effort to do so. The aim of this research was to create a set of improved allometric equations relating the above and belowground biomass to the full range of natural size and growth patterns observed. Four variables (DBH, stem length, small branch number and branch number ≥ 2cm diameter) were highly significant predictors of stem biomass (N≤ 278, p< 0.0001 for all predictors, *complete model* R^2^=0.93). A secondary *field model* (containing DBH and branch number > 2cm diameter), proved highly significant as well (N= 278, p< 0.0001 for both predictors, R^2^=0.84). The belowground biomass was estimated to be 19.2±6.2% of the total dry biomass of the bamboo species examined. To demonstrate the utility of these models in the field and derive stand-level estimates of bamboo biomass, ten 0.36-ha plots were analyzed (N= 3,966 culms), yielding above + belowground biomass values ranging from 4.3–14.5 Mg·ha^-1^. The results of this research provide novel allometric models and estimates of the contribution of *G. weberbaueri* and *G. sarcocarpa* to the total carbon budget of this vast and largely unexplored Amazonian habitat.

## Introduction

### 1.1

The Amazon Basin is a critical component of the global carbon cycle, holding upwards of 100 billion tons of carbon, and will figure prominently into any finance program aimed at curbing anthropogenic carbon emissions and climate change (e.g. Reducing Emission from Deforestation and Degradation – REDD+) (Davidson and others 2012). Within its boundaries, bamboo-dominated forests form a virtually unknown biome the size of all of the primary and secondary forests in Central America combined (UNFAO 2005b). Known as *pacales* in Spanish or *tabocais* in Portuguese, they encompass an estimated 160,000-180,000 km^2^ of tropical lowland forest centered in Southeastern Peru and Southwestern Brazil (Figure 1 inset) (Nelson and others 1997; Smith and Nelson 2011; Carvalho and others 2013). Containing a mixture of trees and two woody bamboo species, *Guadua weberbaueri* Pilger and *G. sarcocarpa* Londoño and Peterson, phylogenetic and fossilized evidence point to the long history of this native habitat (estimated age between 3.12 million – 46,000 years old) (Conover 1994; Olivier and others 2009). Interestingly, the distributional pattern of these forests appear unrelated to any topographic, edaphic or anthropogenic features, save for being broadly located along the Fitzcarrald Arch, a 400,000 km^2^ region of rapid tectonic uplift and weathering between the Andes and the Eastern Amazon Basin (Silman and others 2003; Regard and others 2009). To date, the bamboo-dominated forests remain poorly understood as their existence is often regarded incorrectly as a relic of past disturbance (Griscom and Ashton 2006).

**Figure 1.**
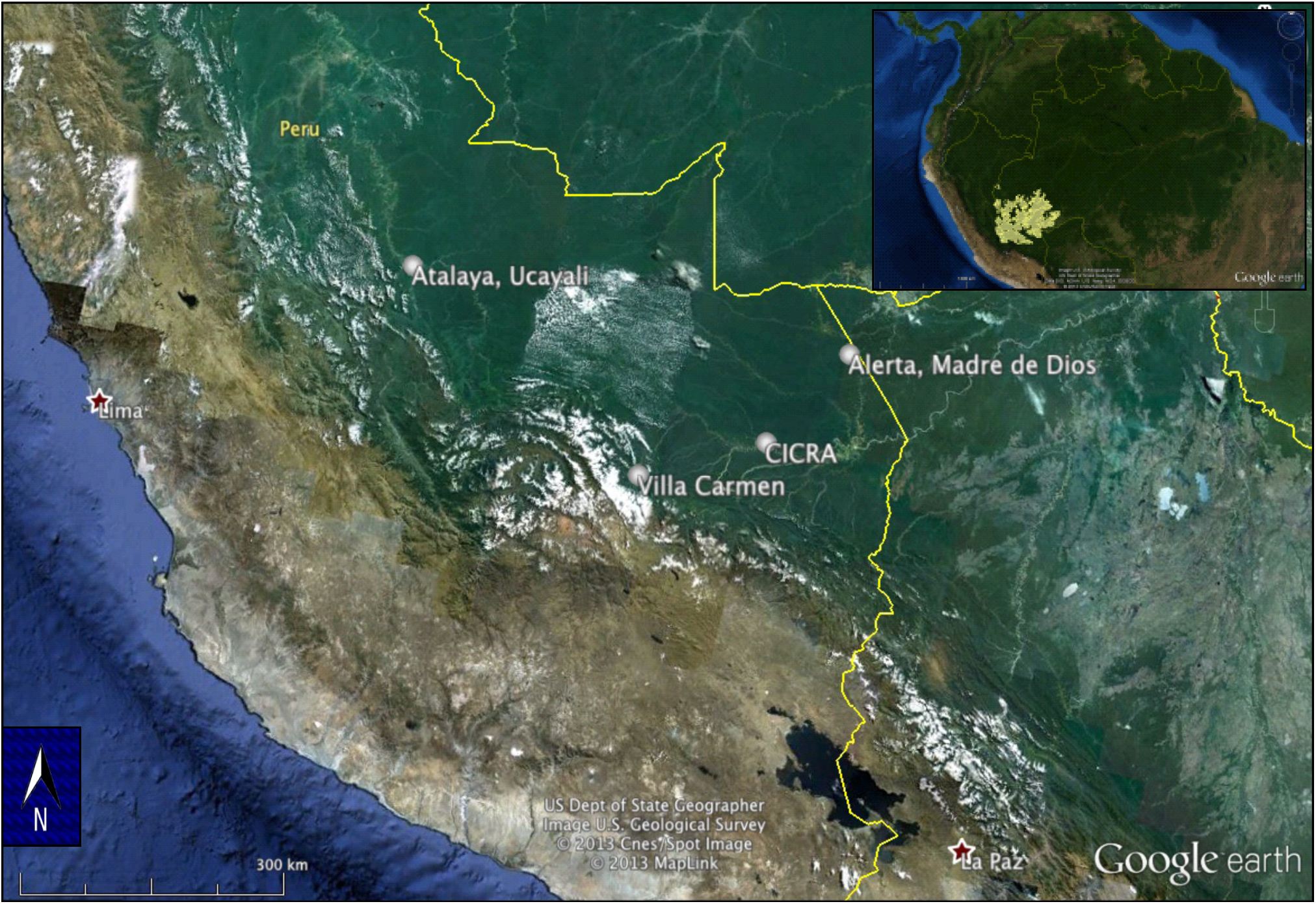
Locations of four sampling sites in SE Peru. (Inset) Estimated 180,000 km^2^ distribution of bamboo-dominated forest (yellow) across SW Amazonia (Nelson and others 1997).

This study sought to improve aboveground allometric relationships for *G. weberbaueri* and *G. sarcocarpa* by encompassing the full range of naturally observed sizes and growth patterns, as well as provide the first estimates of belowground biomass ratios for these two species. Additionally, we strove to develop and implement an easily applicable method of estimating bamboo biomass in the field, with the ultimate goal of highlighting the ranges of bamboo biomass observed across this highly variable habitat. The heterogeneous composition of these forests is well documented, and the internal variability in tree and bamboo biomass, both within (temporal) and between (spatial) individual patches, could have dramatic effects on carbon storage at the landscape and regional scale. Three times more basal area variability in adult trees has been observed inside of bamboo-dominated forests (B+) than in bamboo-free, *terra firme* forests (B-), complicating any comprehensive estimates of the carbon storage of this habitat to date (Silman and others 2003). Further analyses have found that a combination of factors– fewer trees per hectare, decreased wood density of the trees present and decreased tree height resulting from the clambering bamboo– are thought to decrease aboveground biomass by anywhere from 9-56% inside of the B+ forests compared to B-forests (França 2002; Silman and others 2003; Nelson and others 2006; Nogueira and others 2007; Saatchi and others 2007; Nogueira and others 2008; Salimon and others 2011). This range of uncertainty alone is greater than the estimated aboveground biomass for the whole of Canada (Kitani and Hall 1989{Marklund, 2005 #186)}.

Decreasing the level of uncertainty surrounding these values is important for closing the Amazonian forest carbon budget and improving carbon accounting in Southwestern Amazonia (Griscom and Ashton 2006). Presently, the existence of multiple REDD+ projects within the bamboo-dominated forests demonstrates the need for an efficient and accurate method of estimating the contribution of these species to the overall biomass of the region (v-c-s.org 2011). Furthermore, bamboo’s use as a renewable resource for carbon sequestration, timber, pulp, and fiber is becoming an increased topic of research, and quantifying the potential of the Amazonian bamboo-dominated forests for these uses is of significance to various forestry conservation efforts (UNFAO 2005a; Osorio and others 2007; Zhou and others 2011).

### 1.2 Background & Growth

There are an estimated 24 bamboo species of the genus *Guadua* Kunth, the most widespread bamboo genus in the New World, ranging from Mexico to Argentina (Londoño and Peterson 1991). All woody bamboos (Order Poales, Family Poaceae, Tribe Bambuseae) are characterized by complex branching patterns along the main stem (i.e. culm) and two distinct growth phases: shoot elongation and vegetative branching (McClure 1973; Londoño and Peterson 1991). All *Guadua* species, however, are monocarpic perennials that undergo six life stages: flowering, mortality, regeneration, growth, establishment and dominance (Judziewicz and others 1999; Silveira 1999). The two bamboo species present in the B+ forests of SW Amazonia (*G.weberbaueri* and *G. sarcocarpa)* are virtually indistinguishable from each other outside of flowering events as they share ten morphological traits distinct from the remainder of the genus (Londoño and Peterson 1991; Griscom and Ashton 2003).

Growth rates of *G. weberbaueri* and *G. sarcocarpa* can reach 15 cm/day and are characterized by vertical growth for *ca.* 8-12 m before becoming scandent, exploiting nearby trees of similar height and relying on them for support (maximum sizes observed: ca.10.0 cm diameter at breast height [DBH] and > 25 m in height) (Judziewicz and others 1999; Silveira 1999; Nelson and others 2001; Carvalho 2009). This clambering growth pattern has been posited to actively suppress individual tree stems between 5-29 cm DBH (a hypothesis known as *bamboo loading*), and trees greater than this threshold appear to escape any bamboo-derived pressure (Griscom and Ashton 2006). Notably, research from Sena Madureira, Acre, Brazil showed that trees (dicots and palms) ≥ 30 cm DBH maintained 33% lower basal area in B+ forests compared to B-, again pointing to the difficulty in establishing overarching patterns within this diverse habitat (Oliveira 2000). Both *Guadua* species also have the capacity to maintain large quantities of water in their internodes, serving as a habitat for numerous macro- and micro fauna and possibly aiding in their weight-driven suppression of tree stems or offering increased drought tolerance (Louton and others 1996; Vidalenc 2000).

Bamboos propagate vegetatively by underground rhizomes from which all individual ramets of a single genetic individual (i.e. genet) emerge. Rhizome morphology can be broadly split into two parts: the *rhizome proper,* containing all roots and buds (i.e. trunks) from which individual ramets emerge, and *rhizome necks*, thinner portions lacking buds and roots, but responsible for running belowground and the ultimate physical placement of each ramet (Appendix Image 1). In bamboo physiology, roots are a distinctive feature from rhizomes, responsible for the uptake of water and nutrients but comprising a much smaller percentage of belowground biomass (Judziewicz and others 1999). Both the rhizomes and roots of the *Guadua* species analyzed here are largely confined to the upper 50 cm of soil, and their growth strategies differ from many other bamboo genera and congenerics (Silveira 1999). Their phalanx growth strategy allows them to exploit light gaps in the canopy with dense, clumped growth of individual ramets (ca. 30 cm separating individual ramets of a genet). Complementarily, the ability to utilize distant light gaps, termed the guerilla growth strategy, has been documented in these species as well, with rhizomes running underground for distances up to ca. 8 m between ramets (Doust 1981; Smith 2000; Smith and Nelson 2011). This growth pattern greatly enhances lateral mobility at the expense of ultimate rhizome and ramet size (Judziewicz and others 1999). However, the presence of both growth strategies within a single genet, in concert with rapid and suppressive aboveground growth, highlights the phenotypic plasticity of these species and their ability to quickly establish and maintain dominance in these forests (Silveira 1999). Notably, genets have been found to have maximum rhizomal lengths on the order of tens of meters (maximum observed horizontal length was > 30 m in a 10 year old individual), negating the possibility that any significant area is dominated by a single genet (i.e. clone) (Smith 2000).

A distinct property of these bamboo forests is that they occur as large, adjacent patches, each occupied by a single, temporally synchronized cohort derived from mass flowering of the previous monocarpic generation (average life cycle length ca. 28 years, average cohort size ca. 330 km^2^, ca. 480 total cohorts) (Carvalho and others 2013). For the first 8 – 14 years following mass mortality, the majority of the ramets at any one site are juveniles confined to the understory. This lifecycle pattern drives both regional scale variability in bamboo biomass, as well as temporal variability within each patch. However, the total bamboo biomass over the entire range of B+ forests may be relatively constant over time (Carvalho and others 2013).

### 1.3 Previous Allometric Analyses

While greater attention has been focused on the allometric relationship of Asian bamboos, to date, a single previous effort to assess the allometric biomass of *G. weberbaueri* has been carried out (Jyoti Nath and others 2009{Isagi, 1997 #49)}(Torezan and Silveira 2000; Kumar and others 2005). The authors derived a pair of allometric equations from the destructive sampling of ramets in a forest reserve near Rio Branco, Acre, Brazil. Ten individuals comprised of newly emerged shoots and young ramets with no, or few, leafy branches (DBH range from 3.6 – 5.4 cm). A separate group of ten mature ramets with many branches represented a larger size class (DBH range from 4.2 – 5.5 cm). Separate allometries were developed for each group, relating DBH to aboveground biomass. Notably, 96.5% of the ramets in their 3000 m^2^ research area fell within these two size classes. The authors proposed that a 3^rd^ degree polynomial model and a linear model (R^2^ = 0.81 and 0.73, respectively) most accurately predicted biomass of mature bamboo ramets (Appendix Formulae 1a & 1b).

To date, there are no published efforts directed at quantifying belowground biomass (rhizomes and roots) of the bamboo species of interest, even though their potential contribution can be a significant proportion of overall biomass (Riano and others 2002). Please note that throughout this manuscript, it is not our intention to diminish the value of this previous work; however, we sought to build upon their contributions by expanding the range of naturally occurring size classes included, in addition to estimating the fractional contribution belowground biomass. Additionally, the research presented here represents the first published effort to establish any allometric relationship for *G. sarcocarpa*.

### 1.4 Application & Biomass Estimation

Recently, no fewer than three REDD+ projects have been initiated within the bamboo-dominated forests of SE Peru and SW Brazil, with at least two more in the preliminary stages of analysis (v-c-s.org 2011). The three currently existing projects encompasses ca. 90,000 ha of B+ forests (representing ca. 0.5 – 0.6% of the total bamboo distribution) (Nelson and others 1997; Josse and others 2007; Carvalho and others 2013). The methods employed for quantifying the bamboo biomass in these projects has ranged from using a congeneric species of bamboo (*G, angustifolia* Kunth), to conservatively ignoring bamboo’s contribution all together. Thus, even in REDD+’s functional infancy, the importance of accurately quantifying the contribution of these two *Guadua* species is evident.

As a result, the final analyses performed here consisted of measuring all ramets within 3.6-ha of mature B+ forests in accordance with our *field model* derived below. This effort was driven by the objectives of quantifying three remaining unknowns: a) the range of ramet DBH size distribution observed in a natural, mature bamboo forest, b) the natural variability of local bamboo biomass as a function of ramet density, and c) the predictive ability of the allometric equations derived here in comparison to other methods presently employed in estimating the biomass of *G. weberbaueri* and *G. sarcocarpa*.

## Methods

### 2.1 Study Sites

The field component related to the allometric derivation of above and belowground biomass occurred in three locations across southeastern Peru. Preliminary data was collected at the Centro de Investigación y Capacitación Río Los Amigos (CICRA) within the Los Amigos Conservation Concession (Amazon Conservation Association [ACA]; 12°34’9.54” S, 70°6’0.84” W, Figure 1). Supporting data were collected at two separate locations: Hacienda Villa Carmen, managed by ACA (12°53’41.22” S, 71°24’14.88” W) near the town of Pilcopata, Madre de Dios, Peru, and northeast of the city of Atalaya, Ucayali, Peru (10°35’11.52” S, 73°14’34.68” W). To assess the range of ramet size classes observed, and estimate the variability of bamboo biomass contribution in a natural, mature B+ forest, ten 60 x 60 m (0.36-ha) plots were established ca. 10 km NW of the town of Alerta, Madre de Dios, Peru (11°40’27.43” S, 69°18’15.73” W; Figure 1). All of these sites are located on alluvium soils recently deposited (over the last ca. 25 my) from the nearby Andes mountain range, with a mean annual temperature ca. 20°C and mean precipitation ca. 2,500-3,500 mm yr^-1^.

### 2.2 Allometric Aboveground Biomass

In the austral dry season of 2010, 235 ramets inside of CICRA were measured for their diameter at breast height (DBH, 1.3m aboveground), cut at ground level and their culm lengths, branch number < 2 cm diameter (hereafter referred to as *small branch number*) and the number of branches ≥ 2 cm diameter were recorded. All branches were removed and fresh biomass weights were recorded for individual (a) culms, (b) branches + leaves, and (c) all branches ≥ 2 cm diameter + associated leaves. Samples of branches, leaves and culms were dried at 70°C for ca. 48 hours and reweighed to acquire a dry biomass fraction and ultimately correlate dry-to-fresh biomass ratios. In 2011, 33 additional ramets in Hacienda Villa Carmen were measured in an identical manner. Additionally, their corresponding number and percentage of internodes containing water, and culm wall thickness at 1.3 m height were recorded as well. Lastly, in 2012, ten ramets evenly dispersed across a range of DBH sizes ca. 3.5 – 7.5 cm were analyzed in a relatively distant location near Atalaya, Ucayali, Peru to test for allometric consistency with the other sampling locations (N = 278 for DBH and branch # ≥ 2 cm diameter, N = 192 for all other variables mentioned save for internodal water [N = 32] and culm wall thickness [N = 30]). All ramets harvested showed no signs of desiccation or mortality. Primary harvest data is available from the authors upon request.

#### 2.2.1

Allometric equations relating aboveground bamboo biomass to the measured morphological variables were formulated stepwise using least squares fit linear regression models with multiple predictors (*JMP* Version 7.0). An initial *complete model* was explored containing all potential variables: DBH, culm length, small branch number, branch number ≥ 2 cm diameter, number and percentage of internodes containing water, and culm wall thickness. A secondary *field model* was derived containing only those variables readily measurable in the field, DBH and branch number ≥ 2 cm diameter. The latter of these variables is easily characterized by manually shaking the culm and visually counting the number of large branches emerging from the main stem. Such a model is necessary for rapid and easily repeatable estimates of bamboo biomass on the ground. The best models were determined using a stepwise analysis in order to maximize predictive ability and minimize the number of variables included. Lastly, pairwise correlations between all variables were explored for any significant trends and a Pearson’s R value of **XX** was used as a cutoff for significant collinearity.

### 2.3 Belowground Biomass

Eleven 1 x 1m plots were established in 2010 in CICRA, strategically placed in locations containing between 1-4 aboveground ramets in order to encapsulate a wider range of aboveground biomass. All belowground rhizomes and roots were excavated, rinsed, allowed to air dry and weighed. Subsamples of the rhizomes and roots were dried at 70°C for ca. 48 hours and reweighed to acquire a dry-to-fresh biomass constant. Aboveground biomass within these plots was recorded in an identical manner as described in Section 2.2 to correlate the ratio of aboveground to belowground biomass. The fraction of the average contribution of belowground biomass was thus included in our total biomass assessment.

### 2.4 Implementation and Comprehensive Analyses

Ten 0.36-ha plots were established ca. 10 km NW of Alerta, Peru, with ≥ 500 m distance separating each site. Each plot was strategically placed in regions of relatively homogeneous, within-plot ramet densities. Sites representing subjectively low, medium and high ramet densities were prioritized. Within each plot, all ramets were counted and measured for DBH and branch number ≥ 2 cm diameter. A non-parametric (kernel) distribution was fit to the full range of DBH sizes observed and compared to the size classes utilized in the creation of the models derived here as well as the previous allometric models (Torezan and Silveira 2000). The total above and belowground biomass contribution of the bamboo in each plot was estimated in accordance with the *field model* established below, and any trends in biomass as a function of ramet density were explored. Lastly, the derived estimates of total bamboo biomass per plot were compared visually to the other presently employed methods of analyzing bamboo biomass.

## Results

### 3.1 Allometric Aboveground Biomass Estimation

The destructively sampled *G. weberbaueri* and *G. sarcocarpa* ramets had DBH values and culm lengths ranging from 1.8 - 8.1 cm and 0.34 – 24.81 m, respectively. The proportion of dry weight remaining after *ca.* 48 hours of drying subsamples of the culm, branches and leaves was 49.7 ± 3.4% of the fresh weight. Although 49 of our sampled ramets were identified as juveniles (as described by Torezan and Silveira [2000]), incorporating them within the more robust size class distribution of mature ramets permitted for the creation of a single, inclusive model.

Although allometric equations are often defined by power or logarithmic functions, in this study it was found a least squares linear regression model provided the best fit of the biomass data observed (Niklas 2004; Chave and others 2005). The stepwise regression model demonstrated that four variables of the c*omplete model* were highly significant predictors of biomass (p < 0.0001 for DBH, branch number ≥ 2 cm diameter, culm length, and small branch number; Formula A, Table 1). There was no correlation between the number and percentage of internodes containing water (p = 0.75 and 0.88 respectively). Furthermore, stem wall thickness examined over a subset of 30 individuals varied by ca. 25% around the circumference of an individual culm (data not presented). Although its predictive ability was significant (p = 0.01), the limited sampling size and large intra-culm variability observed compelled us to omit this measurement from the final formula of biomass prediction. The resulting variables input into the *complete model* accounted for ca. 93.1% of the variation in biomass observed (Formula A, Table 1, Figure 2):

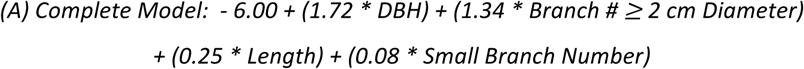

**Figure 2.**
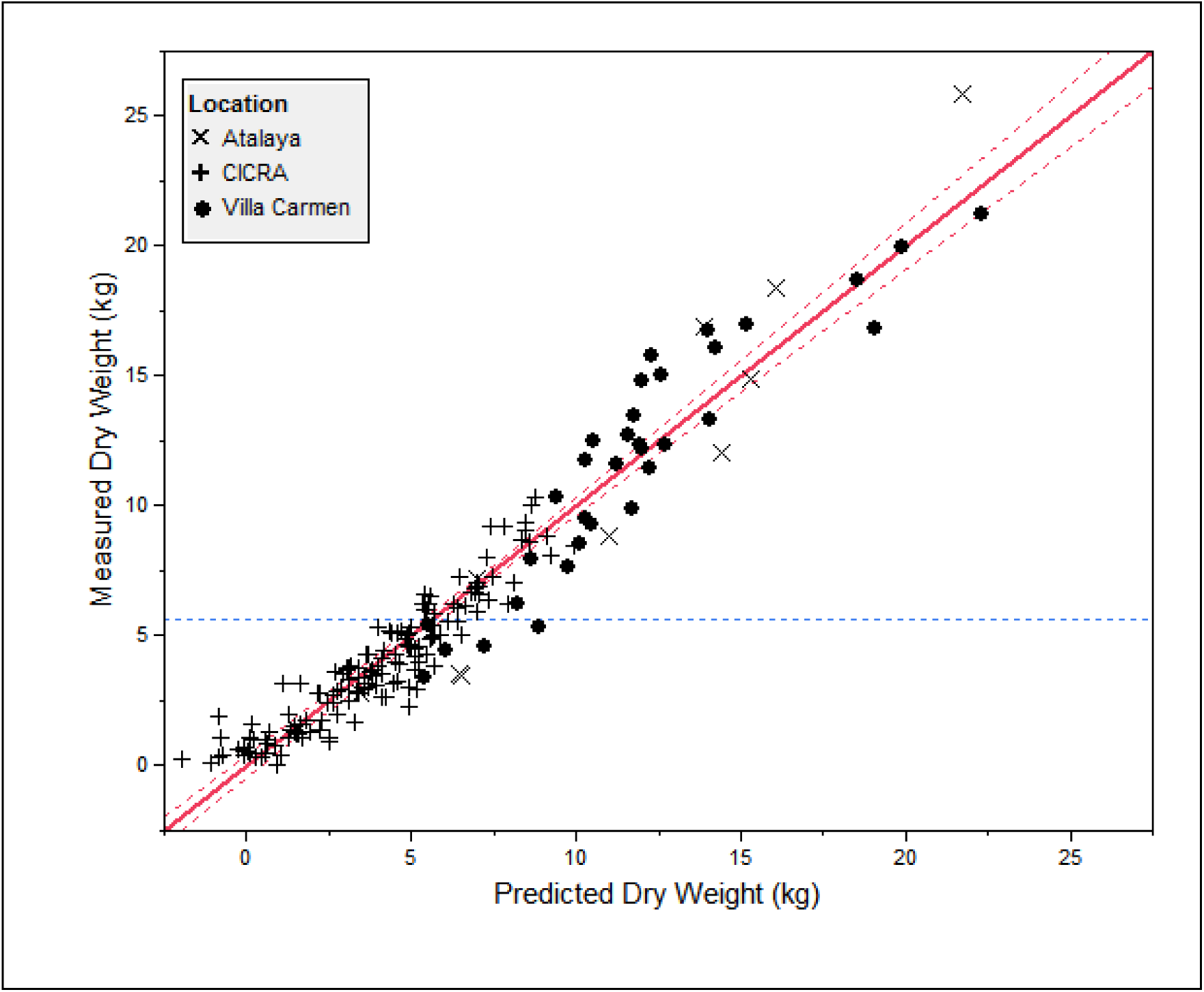
*Complete model* using a least squares fit linear regression and containing the variables DBH, culm length, number of branches ≥ 2 cm diameter and branch number. P < 0.0001 for all variables examined, R^2^ = 0.93, RMSE = 1.24, n = 192. Dashed red lines represent 95% confidence intervals and the horizontal dashed blue line represents the mean weight of all ramets measured.

**Table 1.**

| Parameter | Estimate | Std. Error | P |
| --- | --- | --- | --- |
| *Intercept* | -6.00 | 0.33 | < 0.0001 |
| *DBH* | 1.72 | 0.09 | < 0.0001 |
| *Culm Length* | 0.23 | 0.03 | < 0.0001 |
| *Branch # ≥ 2 cm diameter* | 1.34 | 0.09 | < 0.0001 |
| *Small Branch #* | 0.08 | 0.02 | < 0.0001 |

**Table 2.**

| Parameter | Estimate | Std. Error | p |
| --- | --- | --- | --- |
| *Intercept* | -6.24 | 0.36 | < 0.0001 |
| *DBH* | 2.54 | 0.09 | < 0.0001 |
| *Branch # ≥ 2 cm diameter* | 1.07 | 0.11 | < 0.0001 |

Both variables examined in the *field model* were highly significant predictors as well (p < 0.0001 for DBH and branch number ≥ 2 cm diameter), and accounted for ca. 83.9% of the variation in biomass observed (Formula B, Table 1, Figure 3):

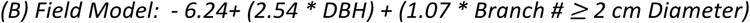

**Figure 3.**
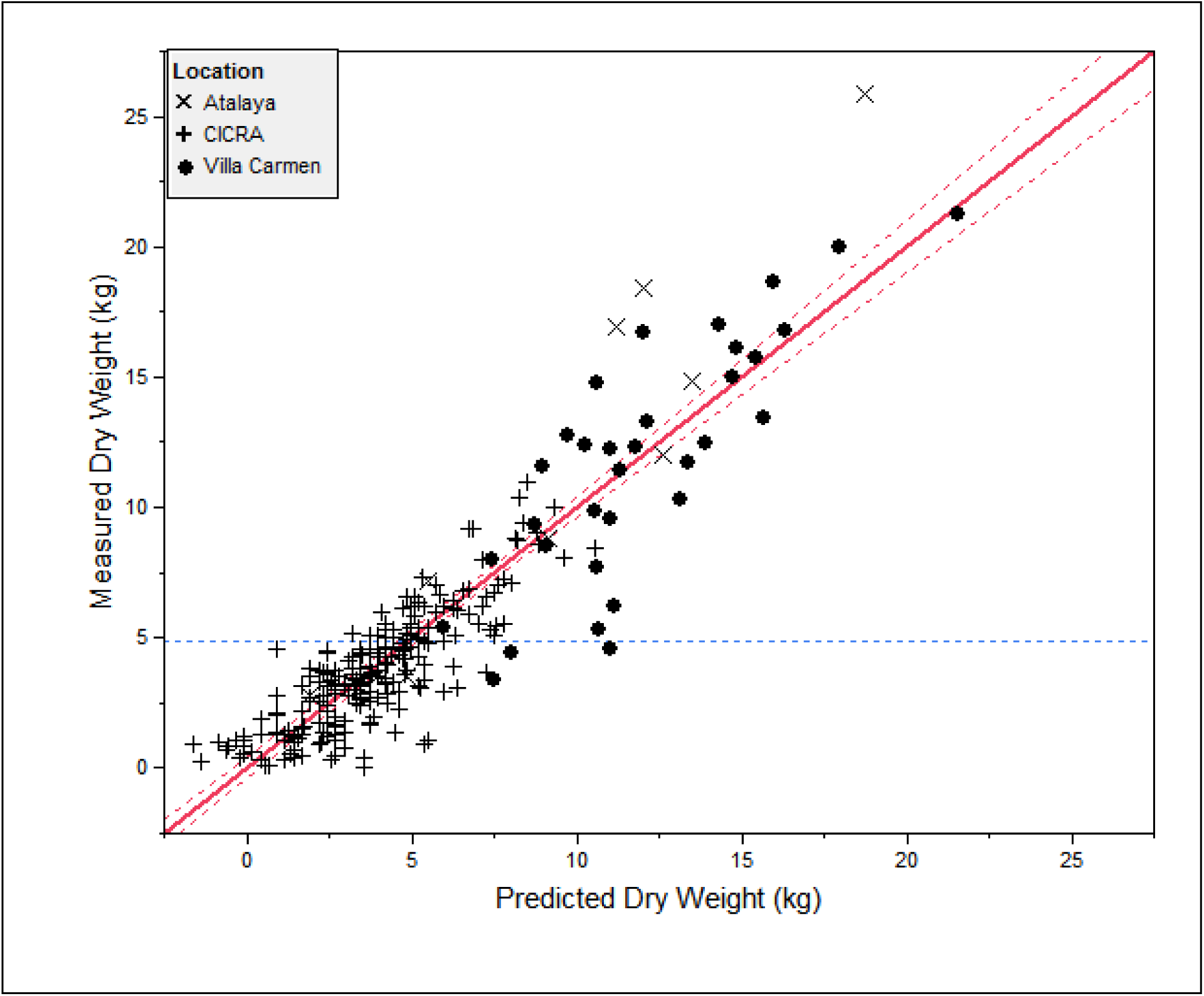
*Field model* using least squares fit linear regression and containing the variables DBH and branch number ≥ 2 cm diameter. P < 0.0001 for all variables, R^2^ = 0.84, RMSE = 1.66, n = 278. Dashed line represents mean weight of all measured ramets. Dashed red lines represent 95% confidence intervals and the horizontal dashed blue line represents the mean weight of all ramets measured.

In particular, DBH alone explained 78.9% of the estimated biomass (Table 1, Appendix Figure 1, Appendix Formula 2). No apparent patterns were observed within the plots of the residuals for each of these three models: *complete, field,* and *DBH-only* (Appendix Figures 2, 3, 4). The Pearson’s R correlations between all model parameters showed that the majority of variables were significantly related to each other (Appendix Table 1). Notably absent from the *field model* variables is *culm length,* as the scandent nature of *G. weberbaueri* and *G. sarcocarpa* often either precludes their apex from sight or their bent culm growth inhibits accurate length estimation. Furthermore, individual ramets were frequently observed to be broken at low heights (< 5 m aboveground, personal observation), diminishing its value as an accurate predictor of aboveground biomass. However, these broken culms were often observed supporting large, thick branches (≥ 2 cm diameter); thus branch number ≥ 2 cm diameter was included in the allometric analyses here.

### 3.2 Belowground Biomass Analysis

The proportion of dry weight remaining of the roots and rhizomes after 48 hours of oven drying was 37.8 ± 19.2% (± 1 S.D.) of the fresh weight. This high variance likely derived from the variation of water content between the two distinct parts of the rhizome: the rhizome proper, containing the buds (ie: trunks) of the ramets, and the long, thin rhizome necks connecting clones of the same individual (see section 1.2, Appendix Image 1). The average belowground fresh weight ratio of the eleven 1 m^2^ plots excavated was 24.0 ± 6.9% of the aboveground fresh biomass. After drying, the average ratio of belowground biomass totaled 19.2 ± 6.2% of the total above and belowground biomass.

### 3.3 Distribution of Biomass

The average percent contribution of each of the morphological features measured (culm, branches + leaves, branches ≥ 2 cm diameter + leaves and rhizome + roots) to total fresh biomass of *G. weberbaueri* and *G. sarcocarpa* across all ramets measured is presented in Figures 4A & 4B for ramets lacking branches ≥ 2 cm diameter (n = 215), and 4C & 4D for ramets with at least one branch ≥ 2 cm diameter (n = 63). Interestingly, the ratio of the contribution of biomass of culms compared to all combined branches (regardless of diameter) and leaves is largely unrelated to culm DBH (R^2^ = 0.03 and 0.15 for ramets without and with branches ≥ 2 cm diameter, respectively). Thus, the analyses here suggest that one would be unable to predict branch + leaf biomass based on DBH size alone.

### 3.4 Natural Size Distribution and Estimated Biomass

Within the ten 60 x 60 m plots (0.36-ha) established in the natural, mature, B+ forest, 3,966 ramets were counted and measured for DBH and branch number ≥ 2 cm diameter. The minimum and maximum ramet densities recorded were 154 and 737 per 0.36-ha, translating to ca. 425 - 2,050 ramets ha^-1^ (ca. 0.04 - 0.2 ramets m^-2^), respectively. The range of ramet diameters observed across all plots was 0.9 – 8.9 cm DBH. By comparison, the range of ramet sizes sampled for the creation of the *complete*, *field* and *DBH-only models* derived here was 1.8 – 8.1 cm DBH, notably short of the observed ranges in the established plots (as well as the documented maximum size of the two species of interest, 10 cm DBH). A kernel probability density distribution was fit to the entire set of data (N = 3,966) and estimated that 0.7% of the ramets measured fell outside of the region used for the calibration of our models here (Appendix Figure 5). The fit density curve also pointed towards a bimodal distribution of size classes, possibly a result of the inherent differences in size growth pattern between *G. weberbaueri* and *G. sarcocarpa (Olivier 2008).*

Plot-level above and belowground bamboo biomass was estimated according to our *field model.* The predicted values of biomass values ranged from 4.3 – 14.5 Mg ha^-1^ (Figure 5 displays aboveground biomass only), with an average above + belowground biomass of 9.7 ± 3.2 kg ramet^-1^ (± 1 S.D.). Median plot DBH values ranged from 4.0 - 6.7 cm DBH (Figure 6). Examining the ramet size distributions showed that the average DBH range observed across all plots was 5.9 ± 1.0 cm (± 1 S.D.). Taking the difference between 5^th^ and 95^th^ % confidence intervals, the range of DBH sizes observed within each plot was 3.0 ± 0.7 cm (± 1 S.D.).

**Figure 4.**
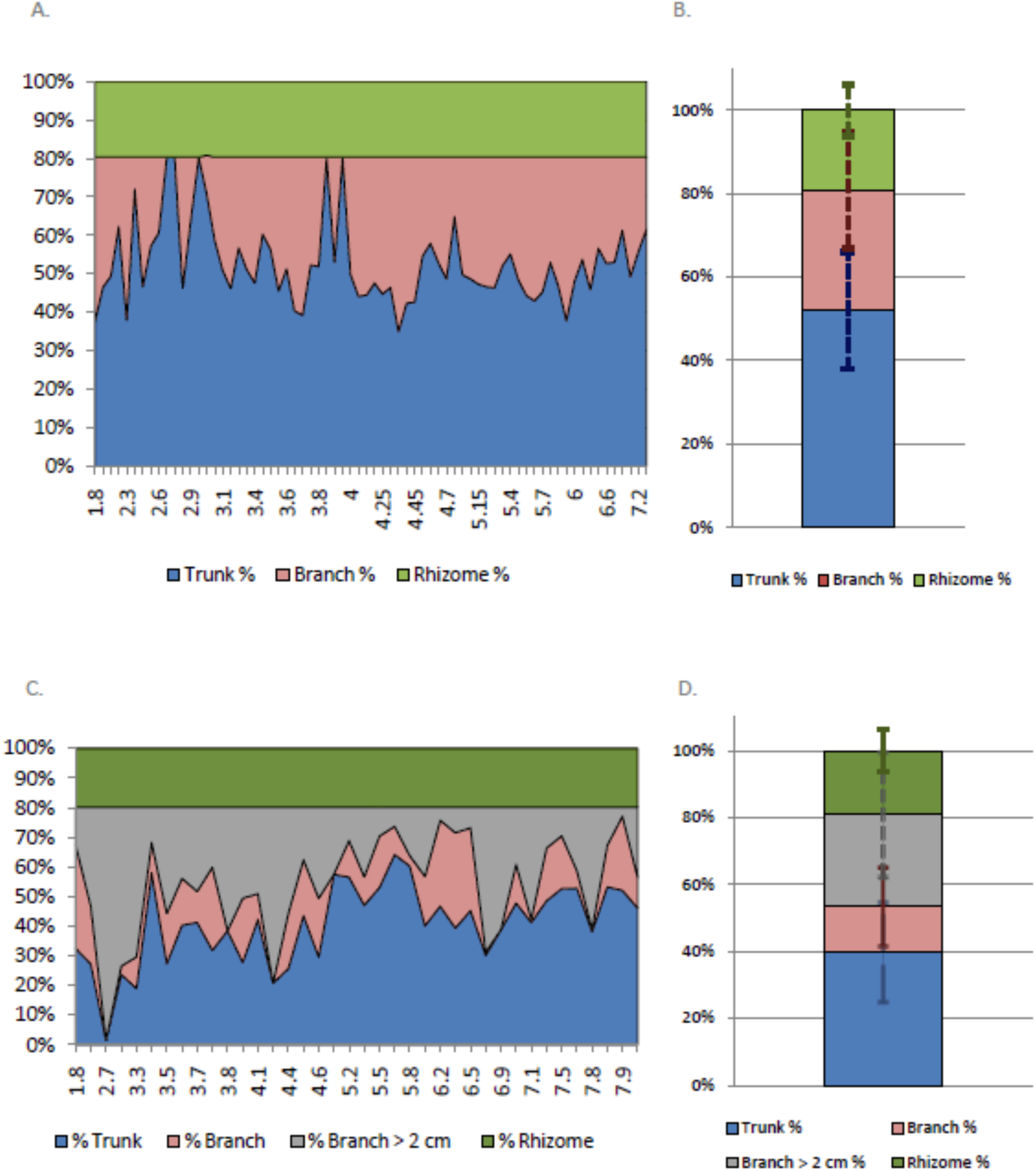
Average fractional contribution of each of the main components measured to the total fresh biomass of each individual by DBH. (A) includes only the average contribution across all individuals without branches ≥ 2 cm diameter (n = 218), and (B) displays the average contribution of biomass (± one standard deviation) across all DBH size classes examined for the same set of ramets. (C) Includes only culms with a at least one branch ≥ 2 cm diameter (n = 60), and (D) displays the average contribution of biomass (± one standard deviation) across all DBH size classes examined for the same set of ramets. In both sets of figures, the percent contribution of rhizomes is fixed at 19.2% ± 6.2 of total above and belowground biomass.

**Figure 5.**
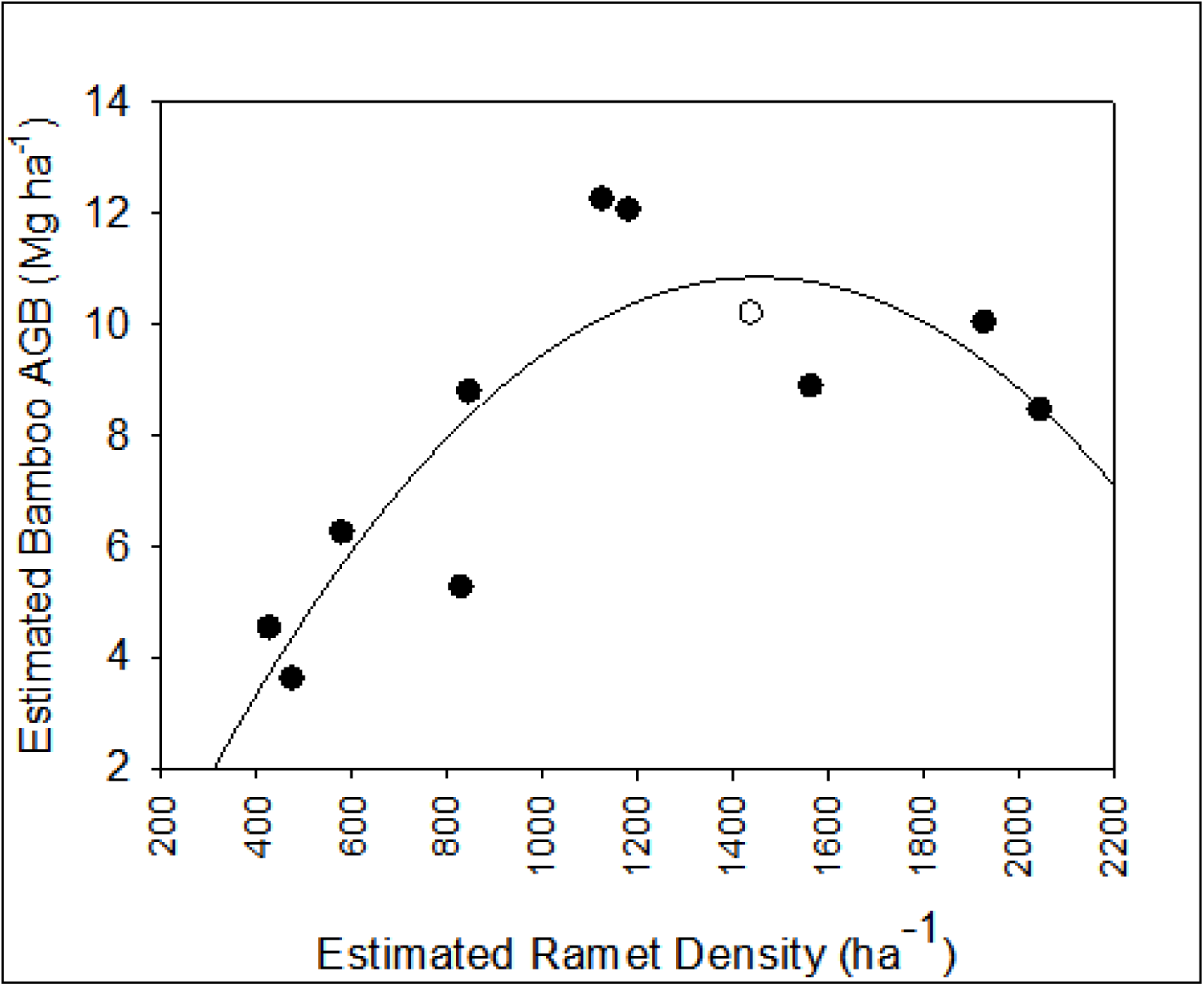
The estimated aboveground biomass contribution across the ten 0.36-ha plots examined (black points). A 2^nd^ degree polynomial equation was selected to fit the observed pattern due to the apparent inflection point at intermediate ramet densities (ca. 1,200 – 1,500 ramets ha^-1^; R^2^ = 0.73 for n = 10). Torezan and Silveira’s (2000) biomass estimate in their 3000 m^2^ assessment of bamboo biomass in Rio Branco, Acre, Brazil is shown (white point, total R^2^ =0.74 for n = 11).

The estimated values of aboveground bamboo biomass correlated with ramet densities across the ten plots examined (Figure 5). At low ramet densities (ca. ≤ 850 ramets ha^-1^), the contribution of bamboo biomass was relatively low (≤ 6.2 Mg ha^-1^). At intermediate ramet densities, (850 < x < 1,200 ramets ha^-1^) bamboo biomass increased to its maximum estimated values (8 < x ≤ 12.5 Mg ha^-1^). Intriguingly, an inflection point appears between. 1,200 – 1,500 ramets ha^-1^, after which the estimated biomass of the bamboo begins to decline (6 ≤ x ≤ 8 Mg ha^-1^; Figure 5). The relationship of mean ramet DBH values as a function of plot ramet density likely explains part of this trend, as mean culm diameters reach their maximum values between 1,200 < x < 1,500 ramets ha^-1^ (Figure 6). Although these estimates represent the first attempts at quantifying the variability of bamboo biomass within the B+ forests, a critical remaining unknown is precisely how the patterns of ramet densities and biomass vary across local, regional and landscape-level scales.

**Figure 6.**
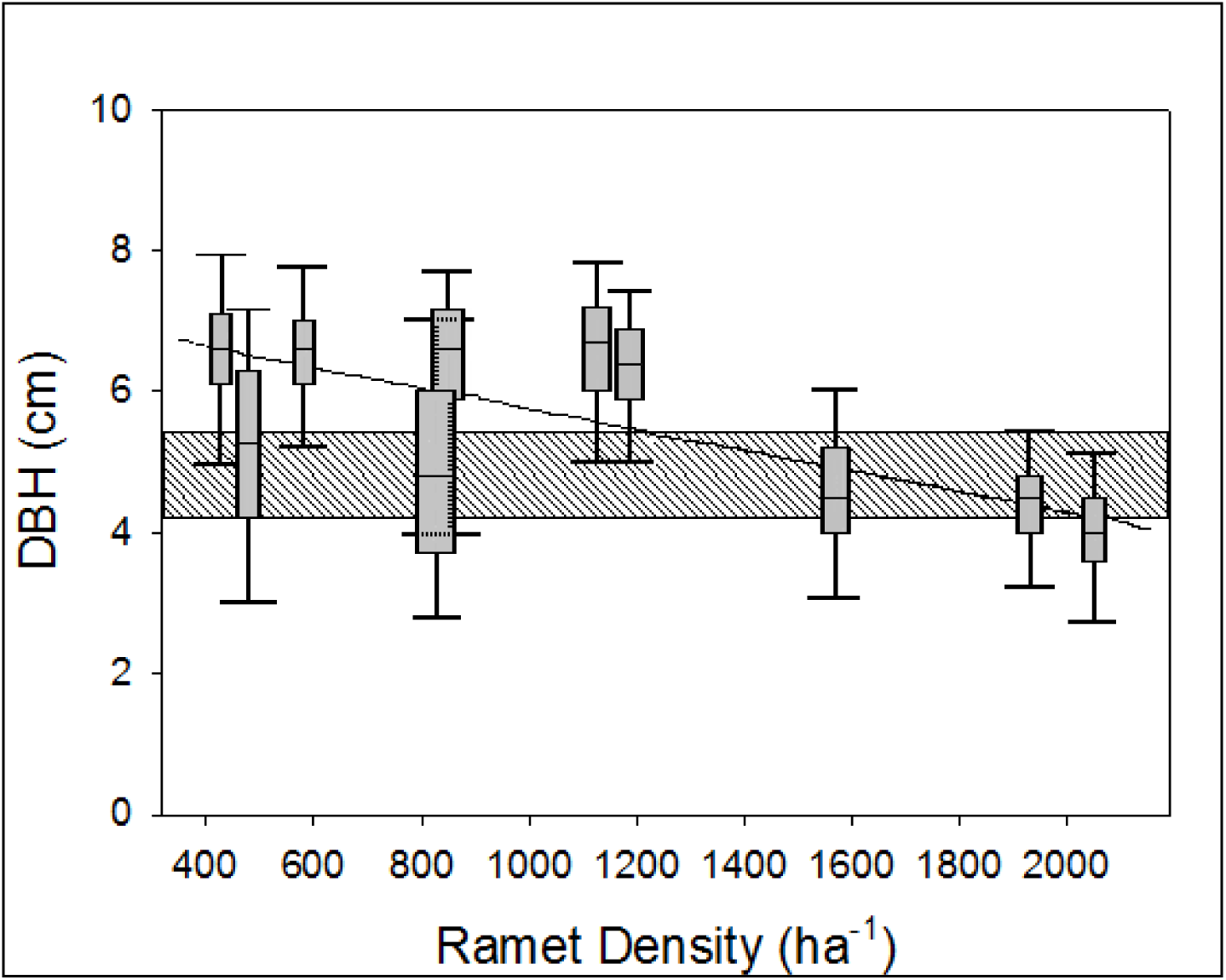
The range of DBH size classes observed within each plot. Lines, boxes, and whiskers represent the median, 50% C.I.’s, and 95% C.I.’s respectively. The range of adult ramets sampled for the previous allometric effort is shown by the grey, hashed rectangle. A simple linear regression was run to characterize the diminishing DBH size distributions as a function of increased ramet densities (hashed line, R^2^ = 0.34)

### 3.5 Comparison to Other Current Methods

The previous allometric models created for one of the two dominant *Guadua* species (*G. weberbaueri*) were based upon the destructive sampling of juvenile (N = 10) and adult (N = 10) ramets in a mature B+ forest near Rio Branco, Acre, Brazil (Torezan and Silveira 2000). The sizes of the ramets sampled ranged from 3.6-5.5 cm DBH (Appendix Equations 1 & 2 refer only to the ten mature ramets sampled, DBH range 4.2-5.5 cm). Again, when taking the difference between the 5^th^ and 95^th^ % confidence intervals, the range of DBH sizes observed within their 3000 m^2^ research area was < 1.9 cm, suggesting their stand was much more homogenous in size distribution than our measured mature B+ forests. When a kernel density distribution was fit to all ramets observed in the 3.6-ha examined (N = 3,966), it was determined that 49.8% of the ramets sampled fell outside of their DBH size classes examined for both mature and juvenile ramets (10.0% of the ramets sampled were < 3.6 cm DBH and 39.8% were > 5.5 cm DBH; Appendix Figure 5).

When compared to our directly measured values of bamboo biomass (N = 278), the fit of their derived linear equation appears to underestimate bamboo biomass at low ramet DBH size classes (< 4.5 cm), and overestimates bamboo biomass for larger sized ramets (> 5.0 cm, Figure 7A). When examining their proposed 3^rd^ degree polynomial equation, the authors’ estimates of biomass become exponentially distorted at DBH size classes < 3.0 and > 6.0 cm (Figure 7B). These figures are presented here solely to represent the fundamental flaw with implementing allometric equations outside of the range over which they were calibrated, a rule well known to Torezan and Silveira (2000) who directly address its importance in their work.

**Figure 7.**
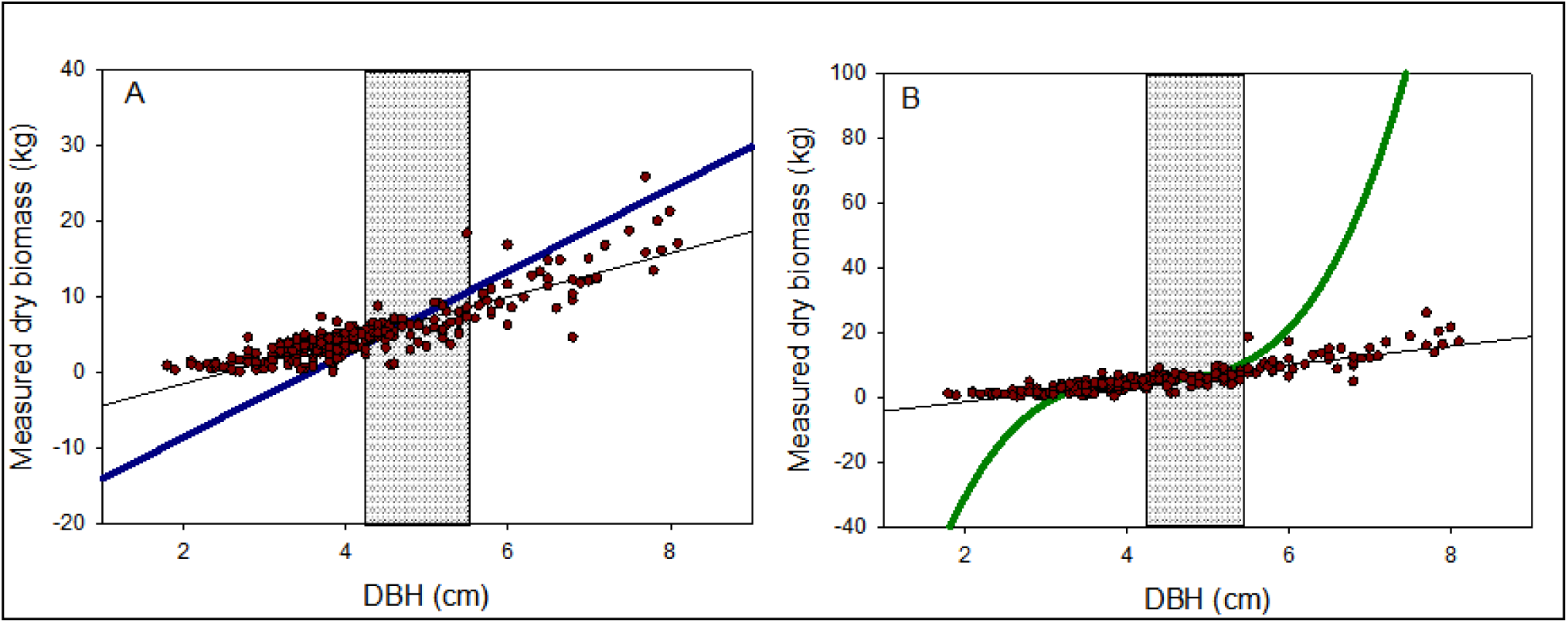
A comparison of the relationships between bamboo DBH (cm), measured dry biomass (kg) and (A) the linear allometric equation proposed by Torezan and Silveira (2000, presented in blue), (B) the third degree polynomial equation proposed by Torezan and Silveira (2000, presented in green). In both figures, the black line represents the *field model* derived in the present research (R^2^ = 0.84, p < 0.0001). The range of DBH sizes (N = 10) used in the creation of their equations is represented by the vertical hashed rectangle.

Torezan and Silveira (2000) implemented their equations across 3000 m^2^ in the same B+ forest from which they sampled their ramets to model bamboo biomass. The authors found 426 ramets within their study site, translating to an average ramet density of 1420 ramets ha^-1^ (0.142 ramets m^-2^). These data are of particular interest as it falls in between the estimated densities of the ten 0.36-ha plots, as well as into the hypothesized zone of inflection (1,200 – 1,500 ramets ha^-1^). The authors estimated the aboveground contribution of bamboo biomass for the entire 3000 m^2^ and derived a value of 10.2 Mg ha^-1^ in this region. When examining this estimate with the data derived from the distinct ten 0.36-ha plots, it appears to fall directly in line with the observed pattern of ramet density and biomass (Figure 5). This not only corroborates the strength of the models created by Torezan and Silveira (2000) for measuring bamboo stands within their size classes examined, but allows for forests separated by > 200 km to support our conclusions regarding the pattern and variability of bamboo biomass in SW Amazonia.

To more readily explore the methods employed by the REDD+ projects established in the region, we made the assumption that the average hectare of B+ forest maintains biomass levels equal to the average of the ten 0.36-ha plots analyzed (above and belowground biomass is 9.5 ± 3.6 Mg ha^-1^). Notably, this value is below the aboveground-only biomass estimate derived by Torezan & Silveira (2000). The 34,000-ha *Purus* project (implemented by *TerraCarbon & CarbonFund*) is located almost entirely within previous estimates of bamboo forest distribution in Acre, Brazil (Nelson and others 1997; v-c-s.org 2011; Carvalho and others 2013). The project developers cautiously chose not to incorporate bamboo into their biomass assessment. Using the average estimate of bamboo biomass ha^-1^, the utilization of the *field allometric model* could potentially add > 300,000 Mg of biomass to their project (comparable to > 1,000-ha of old growth, bamboo-free terra firme forest) (Saatchi and others 2007).While this value must be taken with caution until accurate estimates are obtained regarding bamboo ramet density and biomass patterns across the region, this value still highlights the importance of the contribution of these two bamboo species to such carbon mitigation strategies.

Alternatively, two distinct REDD+ projects cover ca. 38,500-ha of bamboo-dominated forests according to a 2010 vegetation map created by the Peruvian Ministry of the Environment (Ministerio del Ambiente [MINAM], as cited within each report) (v-c-s.org 2011). In each of these projects, the densities of *G. weberbaueri* and *G. sarcocarpa* ramets were sampled and biomass was estimated using the average value for a congeneric species, *G. angustifolia* Kunth, commonly found in northwestern South America (Judziewicz and others 1999). A standard estimate of 47.022 kg ramet^-1^ was applied to all ramets observed. If such a method were applied to each of the ten 0.36-ha plots sampled, the derived estimate of bamboo biomass is consistently an order of magnitude greater than those derived from the *field model* (Figure 8). By extrapolating our average bamboo biomass value throughout the entire distribution of the B+ forests contained within the boundaries of these projects, it is conceivable that the authors overestimated biomass by ca. > 1,500,000 Mg of biomass (as much biomass as in > 5,500-ha of old growth, terra firme B-forest) (Saatchi and others 2007). Again, this value should serve solely as a preliminary estimate until regional patterns of bamboo density and biomass can be established.

**Figure 8.**
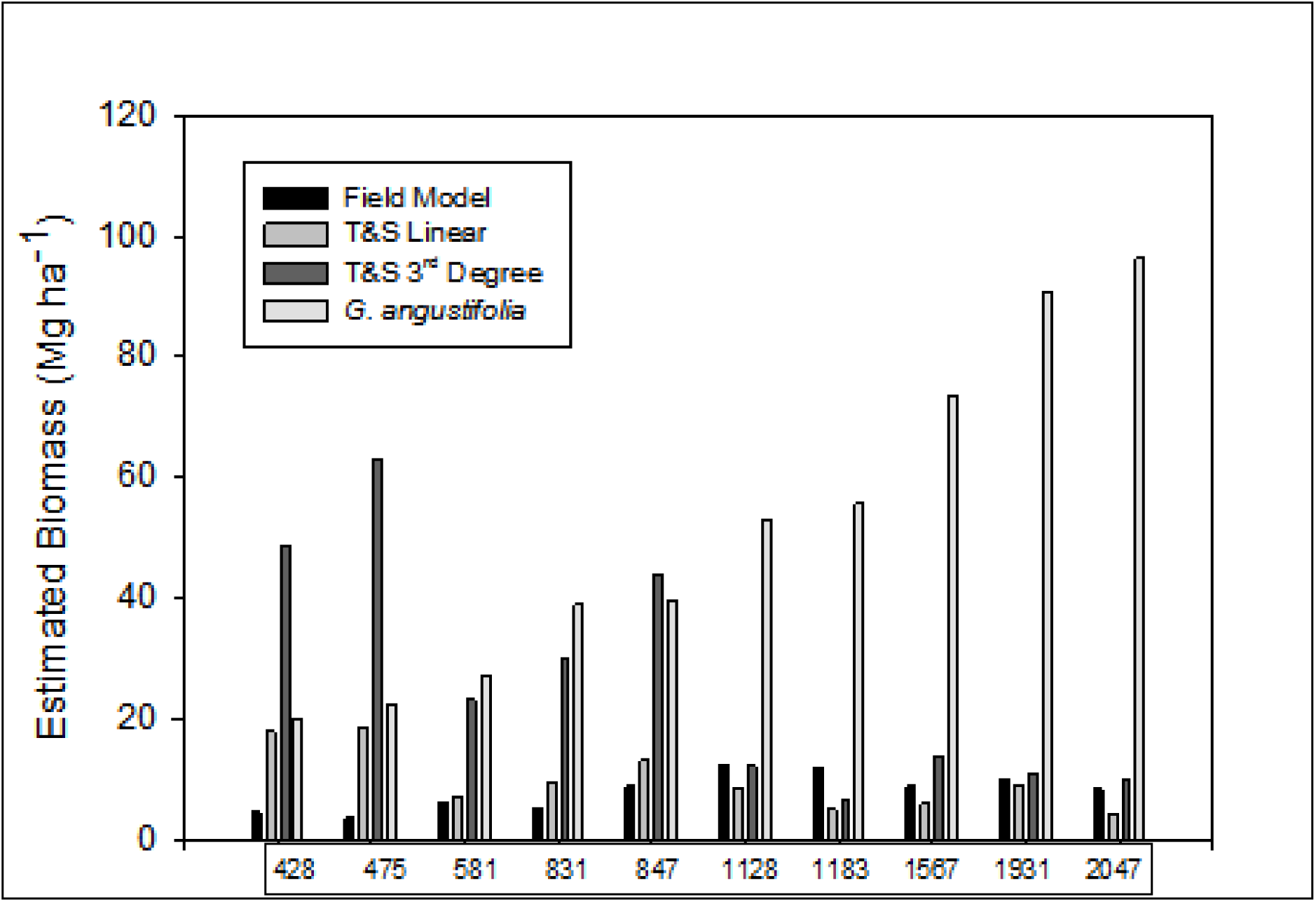
Comparison of the estimated aboveground (only) biomass across the ten 0.36-ha plots using the *field model* derived in the present research, the linear and 3^rd^ degree polynomial allometric models proposed in a previous allometric study (Torezan and Silveira 2000), and the use of a congeneric species (*G. angustifolia)* in two REDD+ projects currently established.

## Discussion

### 4.1

Although *Guadua weberbaueri* and *G. sarcocarpa* appear to contribute only a fraction of the aboveground biomass to the bamboo-dominated forests of SW Amazonia, their dramatic influence on structure, carbon content and dynamics of this significant habitat is well documented, but as of yet, poorly quantified (Torezan and Silveira 2000; Silman and others 2003; Nogueira and others 2008). An important first step in understanding these relationships and determining these species value as a source of carbon sequestration, timber, pulp and fiber, is to establish a rapid and accurate allometric method for biomass estimation. To date, however, the variability surrounding the biomass estimates of these two bamboo species mandated that an updated analysis of bamboo biomass be carried out.

The allometric relationships derived here allow for the accurate estimation of the biomass of the two bamboo species analyzed. The *complete model* explained 93.1% of the variability in aboveground biomass of individual ramets of *G. weberbaueri* and *G. sarcocarpa*. Additionally, the *field model* showed significant potential as a more rapid biomass assessment technique on the ground, explaining 83.9% of the variability measured in the aboveground biomass. The strength of DBH as a useful predictor of aboveground biomass is expected; however, the introduction of the variable *branches ≥ 2 cm diameter* added significantly to the predictive capacity of these models.

The estimated ratio of belowground biomass of *G. weberbaueri* and *G. sarcocarpa*, 19.2 ± 6.2% of total biomass, to the best of the authors’ knowledge, represents the first attempt at quantifying the belowground rhizome and root biomass of these species. The similarity between the belowground biomass estimates acquired here and those of the congeneric species (*G. angustifolia* Kunth, 19.9%) lends support to these findings (Riano and others 2002).

Interestingly, the number and percentage of internodes filled with water were not found to significantly correlate to biomass..The bamboo species examined here are known to store water in intact internodes, a microhabitat for many invertebrates (Louton and others 1996). Further hypotheses address the internodal water’s ability to alter the habitat’s hydrology, potentially increasing these bamboo species tolerance to drought and/or weight-driven suppression of tree growth (known as *bamboo loading,* see Section 1.2) (Vidalenc 2000; Griscom and Ashton 2006). The insignificant relationships found in the complete model between ramet biomass and the number and percentage of internodes containing water is somewhat surprising when considering the potential function of internodal water as a mechanism to increase ramet weight and constrain tree growth. One could readily hypothesize a potential correlation where either larger or smaller culms store greater amounts of water to increase their mass to a critical level capable of damaging tree branches and stems. When directly examining the percentage of internodes containing water as a predictor of aboveground biomass, a weak, although significant, relationship is revealed (R^2^ = 0.14, p = 0.03). However, the ability of these species to opportunistically uptake and retain water as a drought coping mechanism, noting their inability to access deep soil water with all rhizomes and roots existing in the upper 50 cm, could result in the seemingly random distribution of internodal water.

When examining culm length as a predictor of total biomass, it was discovered that often culms were broken off at relatively low heights (< 5 m). Furthermore, these broken culms often displayed large branching (≥ 2 cm diameter) below any damage. Large branching was also common among larger, unbroken culms capable of supporting such branches. Indeed, when examining culm length as a predictor of the total weight of branches ≥ 2 cm diameter (with associated leaves) for all individuals with at least one branch meeting this criteria, there was no significant correlation found (n = 63, R^2^ = 0.02, p = 0.30). Two potential explanations for this observed phenomena are internal, mechanical disturbance caused by constant ramet turnover, or ramet growth reaching unsustainable heights in locations without nearby support for scandent growth. As a result of either scenario, the breakage of the main culm results in a shift in resources towards larger branching of new leader shoots below.

The estimated above+belowground biomass of the N = 3, 966 ramets measured across the ten 0.36-ha plots established near Alerta, Madre de Dios Peru was 4.3 – 14.5 Mg ha^-1^ (varying by > 300%), with an average of 9.7 ± 3.2 kg ramet^-1^ (± 1 S.D.). When a kernel density distribution was fit to the size distribution of the ramets measured, a bimodal pattern emerged, potentially indicative of the difference in growth patterns between the smaller *G. weberbaueri* and larger *G. sarcocarpa (Olivier 2008).* Indeed, the smaller of the two pulses does appear to correlate closely with the size classes examined by Torezan and Silveira (2000) in their allometric assessment of *G. weberbaueri* (Appendix Figure 5). Intriguingly, bamboo stand biomass appeared to correlate with ramet stand density, with low culm densities maintaining relatively low bamboo biomass from the few ramets present, medium culm densities maintaining the highest levels of biomass from the larger average ramet DBH found within these forests, and the highest culm densities maintaining levels of biomass in between the previous two estimates deriving from the decreased average ramet DBH inside of these forests. The hypothesized mechanism that could be driving this observed patterning is that at the highest culm densities, fewer trees are present to permit for the scandent growth of these two bamboo species. As a result, the bamboo becomes locally hyper-abundant, but its ability to reach larger size classes is resource-limited without access to the canopy. Although the estimated contribution of bamboo biomass from a location 200 km away appears to align with this pattern, these findings still require further investigation from other locations within the B+ forest (Torezan and Silveira 2000).

### 4.2

The previous allometric models created by Torezan and Silveira (2000) were based upon the destructive analysis of ten adult ramets (compared to N = 278 in the analyses here), and both of the authors’ proposed equations appear to underestimate aboveground bamboo biomass at lower DBH sizes, and overestimate biomass at greater DBH sizes. Their linear allometric formula does not permit any culm below a DBH of 3.55 cm to contribute to total biomass (ie: all DBH values < 3.55 cm yield a negative biomass estimate in their equation), leading to an underestimate of aboveground biomass at lower DBH size classes (Figure 7a, Figure 8). Additionally, their 3^rd^ degree polynomial equation showed errors in similar directions and are exaggerated exponentially (Figure 7b, Figure 8). Because 96% of the stems in their study area fell within the size classes used for the development of their allometric models, their estimated aboveground biomass of the bamboo can be assumed accurate and correlated very closely with the findings from the separate ten 0.36-ha plots examined here (Figure 5). However, it is likely that their comparatively smaller areal coverage contributed to the limited distribution of DBH size classes observed. Despite the well known warnings against implementing an allometric model for a size class outside which it was calibrated, their linear model was employed to an undisclosed degree in a high profile, analysis of aboveground biomass in SW Amazonia (Torezan and Silveira 2000; Asner and others 2010). From the current authors’ understanding, it was used to estimate the biomass of all ramets ≥ 10 cm DBH, a size far removed from the mature culm calibration values of 4.2-5.5 cm DBH. Although the abundance of ramets in this size class is likely quite rare, the analyses here suggest the potential for significant overestimation of aboveground bamboo biomass for the B+ regions examined in this Amazonian research.

When examining the various REDD+ projects implemented in the bamboo-dominated forests, two projects used a single value derived from a congeneric species, *G. angustifolia,* averaging 47.022 kg ramet^-1^ (Cruz 2009). *G. angustifolia* can reach 25 cm DBH (average, 9-12 cm DBH) and can reach 30 m in height. Our two bamboo species (*G. weberbaueri, G. sarcocarpa)* have a maximum recorded size of 10 cm DBH and 25 m in height (Judziewicz and others 1999). The average DBH from the N = 3,966 ramets measured was 5.6 ± 1.0 cm (± 1 S.D.) and the average biomass was 9.7 ± 3.2 kg ramet^-1^. As a result, the use of *G. angustifolia* as a surrogate for biomass of the other two *Guadua* species of interest does not appear to be an accurate substitution. The REDD+ authors of the two projects implementing this technique further concluded that bamboo-dominated forests maintained the highest levels of biomass of all forest types examined, a finding that is contradictory to nearly all of the published literature on these forests (França 2002; Silman and others 2003; Nelson and others 2006; Nogueira and others 2007; Saatchi and others 2007; Nogueira and others 2008; Salimon and others 2011).

Looking more closely at the derived estimates of plot biomass using the two previously created allometric equations and the substitution of *G. angustifolia*, at low ramet densities (< 1,000 ramets ha^-1^), all three of these models consistently predict elevated levels of biomass compared to the *field model* derived here (Figure 8). For the two allometric models, this is the result of the average ramet DBH size classes observed within these forests being greater than the range for which they were calibrated (Figure 6). At intermediate ramet densities 1,000 < x < 1,500 ramets ha^-1^) and above, the two previous allometric models are more similar to the estimates derived from the field model, a result of the decreasing average ramet DBH with increasing ramet density. However, even these more comparable estimates of biomass varied up to ca. ± 60% from the *field model* derived estimates (Figure 8). The magnitude of error associated with using of *G. angustifolia* as a surrogate for *G. weberbaueri* and *G. sarcocarpa* increases linearly with estimated ramet density (as these biomass estimates come directly from applying an average ramet biomass to all observed ramets within a region). Although REDD+ projects have only begun taking shape in the past couple years, these extreme examples - from conservatively neglecting bamboo biomass, to vastly overestimating the average biomass contribution of individual ramets using unsuitable alternative methods - substantiates the need for an encompassing and rapid method of assessing the biomass contribution of *G. weberbaueri* and *G. sarcocarpa* across this vast region.

## Conclusions

### 5.1

The allometric relationships established here have shown significant potential in estimating the biomass across the range of size classes observed in mature, natural bamboo-dominated forest. Additionally, estimates of the contribution of belowground biomass *G. weberbaueri* and *G. sarcocarpa* provide further insight into characterizing the dynamics and quantifying the carbon content of this largely unknown habitat. Moving forward, these relationships represent a critical first step towards stand-level assessments of biomass inside of the bamboo-dominated forest of SW Amazonia and the elucidation of any patterns relating to bamboo-tree stem densities and biomass. Many important critical unknowns remain, however. Accurately accounting for the biomass of these two bamboo species is just one issue for future and current REDD+ project managers. The importance of incorporating the observed 28 year temporal lifecycle into successful REDD+ projects is mandatory, as entire cohorts (average size 340 km^2^) undergo mass mortality and subsequently lose all of their biomass over the course of just few months to years (Carvalho and others 2013). Finding a method to incorporate this growth pattern, as well as examining any changes in bamboo ramet density or biomass through subsequent lifecycles, is essential to successfully implementing such carbon mitigation schemes within this vast habitat. Furthermore, characterizing the variability of bamboo ramet densities and biomass across regional and landscape-level scales is necessary for accurately estimating the biomass of bamboo for these two species and removing the notion that the 160,000 - 180,000 km^2^ of bamboo-dominated forest is a single, homogeneous entity.

## Acknowledgements

Funding was acquired through the Amazon Conservation Association (ACA-Washington DC), Wake Forest University (WFU) Vecellio Fellowship and the WFU Center for Energy, Environment and Sustainability (CEES). We thank Manu National Park and Servicio Nacional de Áreas Naturales Protegidas por el Estado (SERNANP) of Peru, ACA for their financial aid and infrastructural support in the lowland forests of Peru, the Silman Lab, Rebecca Dickson, Rebecca Powell and Kathryn Riley (WFU) for their support and input. Lastly, we thank the many field assistants who assisted with data collection; notably Luis Ynmunda, Edgard Collado Delgado and Graham McHenry.

## Appendix

**Appendix Table 1.** Pairwise Pearson’s R correlations between all variables examined.

| **Variable** | **by Variable** | **Correlation**<br>**(Pearson's R)** | **N** | **p-Value** |
| --- | --- | --- | --- | --- |
| Culm Length | DBH | 0.62 | 278 | < 0.0001 |
| Branch # ≥ 2 cm | DBH | 0.42 | 278 | < 0.0001 |
| Branch # ≥ 2 cm | Length | 0.19 | 278 | = 0.001 |
| Complete Dry Weight | DBH | 0.89 | 278 | < 0.0001 |
| Complete Dry Weight | Length | 0.74 | 278 | < 0.0001 |
| Complete Dry Weight | Branch# ≥ 2 cm | 0.58 | 278 | < 0.0001 |
| Small Branch Number | DBH | 0.54 | 192 | < 0.0001 |
| Small Branch Number | Length | 0.78 | 192 | < 0.0001 |
| Small Branch Number | Branch# ≥ 2 cm | 0.07 | 192 | = 0.37 |
| Small Branch Number | Complete Dry Weight | 0.63 | 192 | < 0.0001 |
| % Internodes Containing Water | DBH | -0.26 | 32 | = 0.16 |
| % Internodes Containing Water | Length | -0.34 | 32 | = 0.05 |
| % Internodes Containing Water | Branch# ≥ 2 cm | 0.00 | 32 | = 0.99 |
| % Internodes Containing Water | Complete Dry Weight | -0.38 | 32 | = 0.03 |
| % Internodes Containing Water | Small Branch Number | -0.39 | 32 | = 0.03 |
| Culm Wall Thickness | DBH | 0.49 | 30 | = 0.006 |
| Culm Wall Thickness | Length | 0.28 | 30 | = 0.14 |
| Culm Wall Thickness | Branch# ≥ 2 cm | 0.19 | 30 | = 0.31 |
| Culm Wall Thickness | Complete Dry Weight | 0.58 | 30 | < 0.0001 |
| Culm Wall Thickness | Small Branch Number | 0.28 | 30 | = 0.13 |
| Culm Wall Thickness | % Internodes Containing Water | -0.58 | 30 | < 0.0001 |
| # Internodes with Water | DBH | -0.24 | 32 | = 0.18 |
| # Internodes with Water | Culm Length | 0.04 | 32 | = 0.84 |
| # Internodes with Water | Small Branch Number | -0.08 | 32 | = 0.65 |
| # Internodes with Water | Branch # ≥ 2 cm | 0.02 | 32 | = 0.92 |
| # Internodes with Water | Culm Wall Thickness | -0.47 | 30 | = 0.009 |
| # Internodes with Water | % Internodes Containing Water | 0.86 | 32 | < 0.0001 |

**Appendix Image 1.**
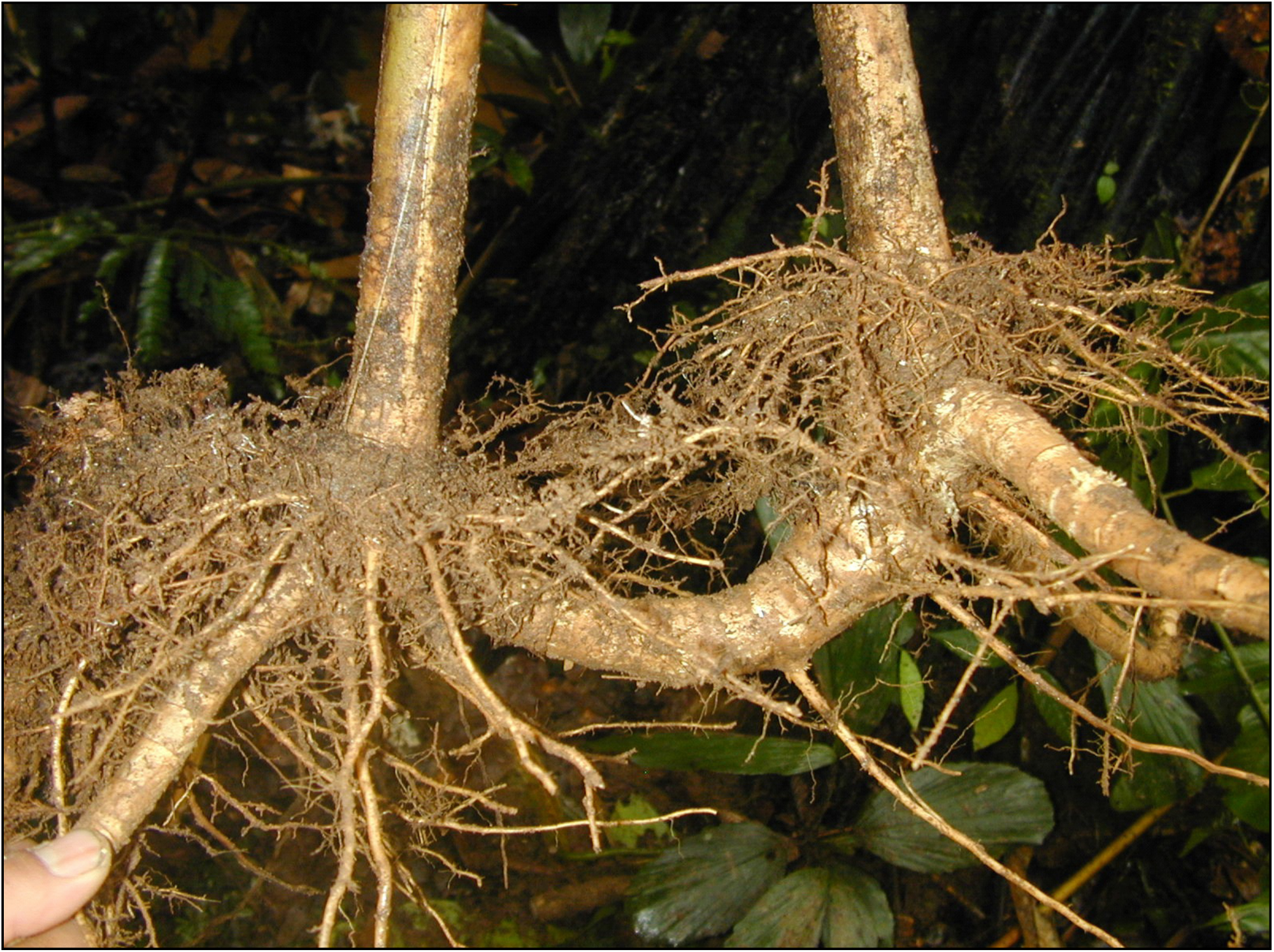
Morphological features of the underground rhizomes of *Guadua weberbaueri* and *G. sarcocarpa*. *Rhizome proper* features are outlined in dashed red circles, while *rhizome necks* are outlined in solid blue ovals (photograph taken by William Farfan Rios).

**Appendix Figure 1.**
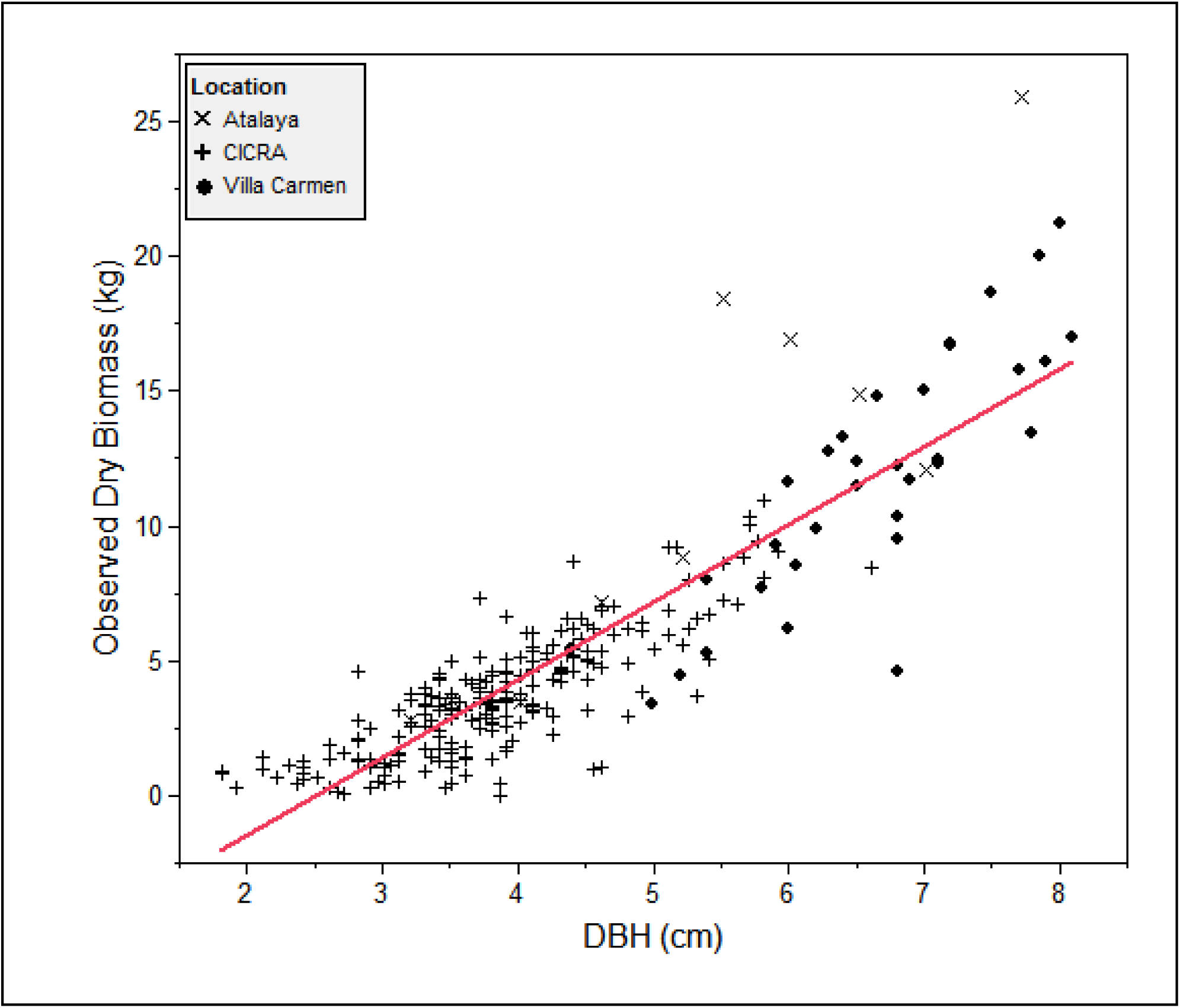
*DBH-only model* using least squares fit linear regression. P < 0.0001 for DBH, Adj. R^2^ = 0.79, RMSE = 1.90, n = 278. Solid red line represents mean weight of all measured ramets.

**Appendix Figure 2.**
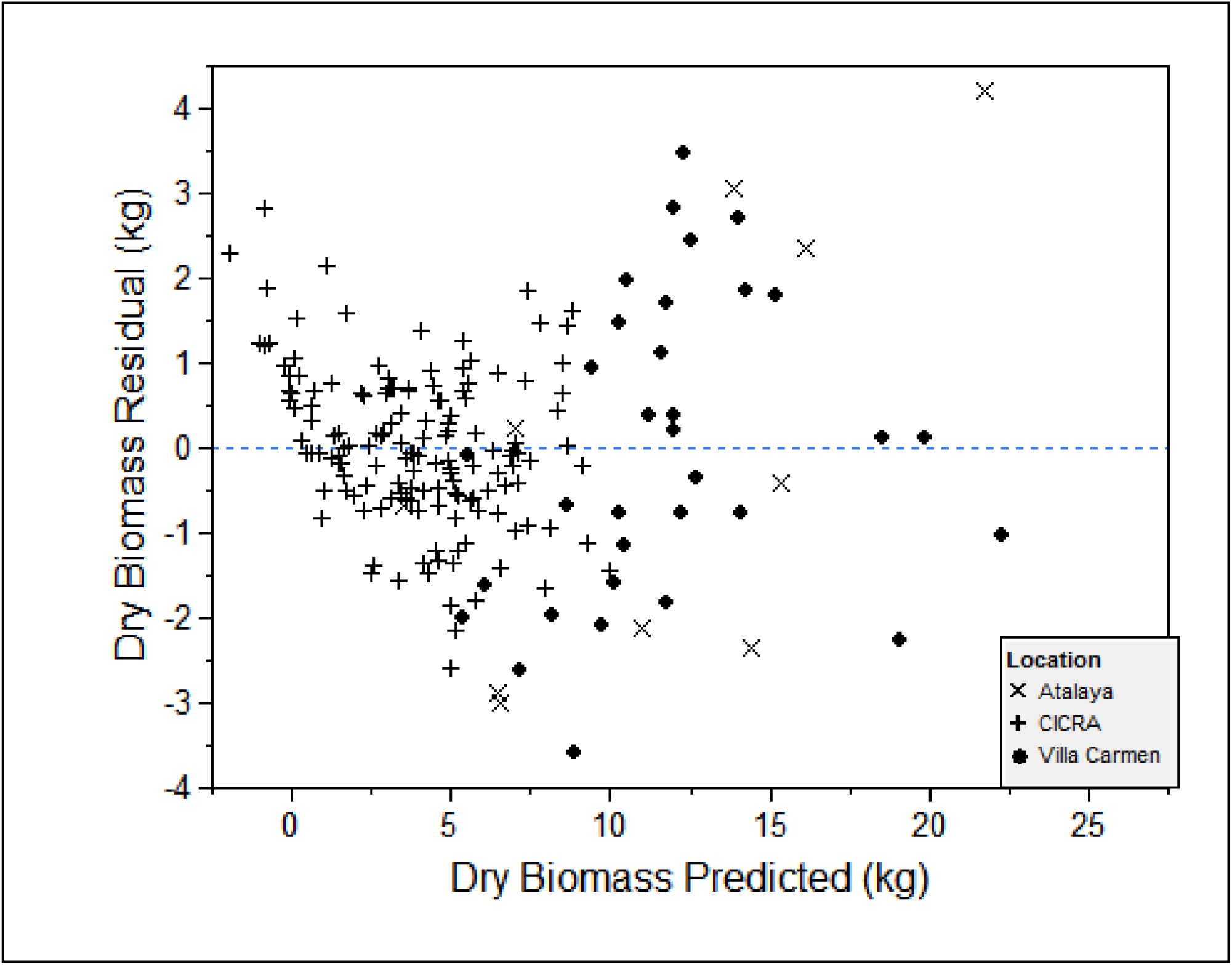
Plot of the predicted residuals for the *complete model.*

**Appendix Figure 3.**
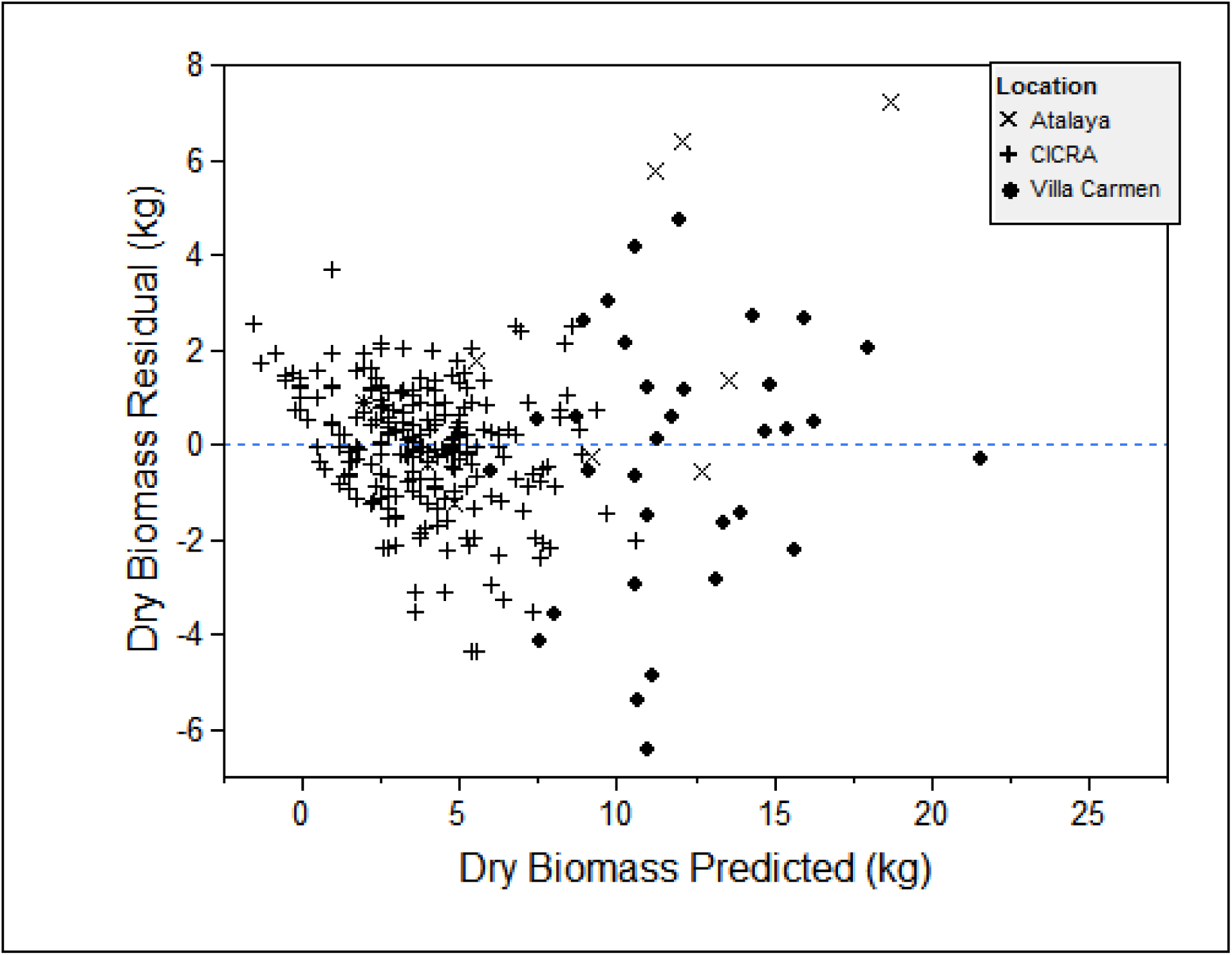
Plot of the predicted residuals for the *field model*

**Appendix Figure 4.**
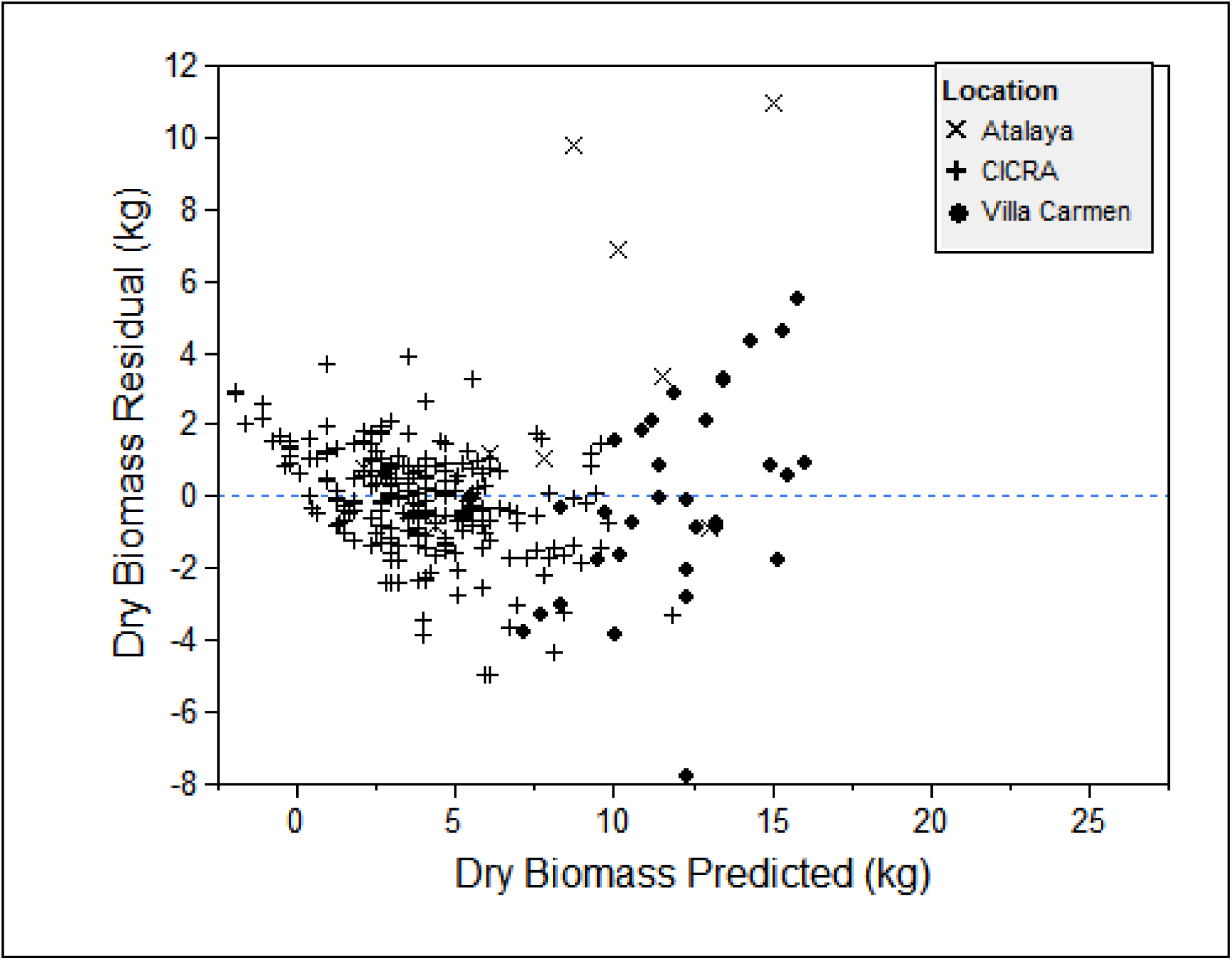
Plot of the predicted residuals for a *DBH-only* model.

**Appendix Figure 5.**
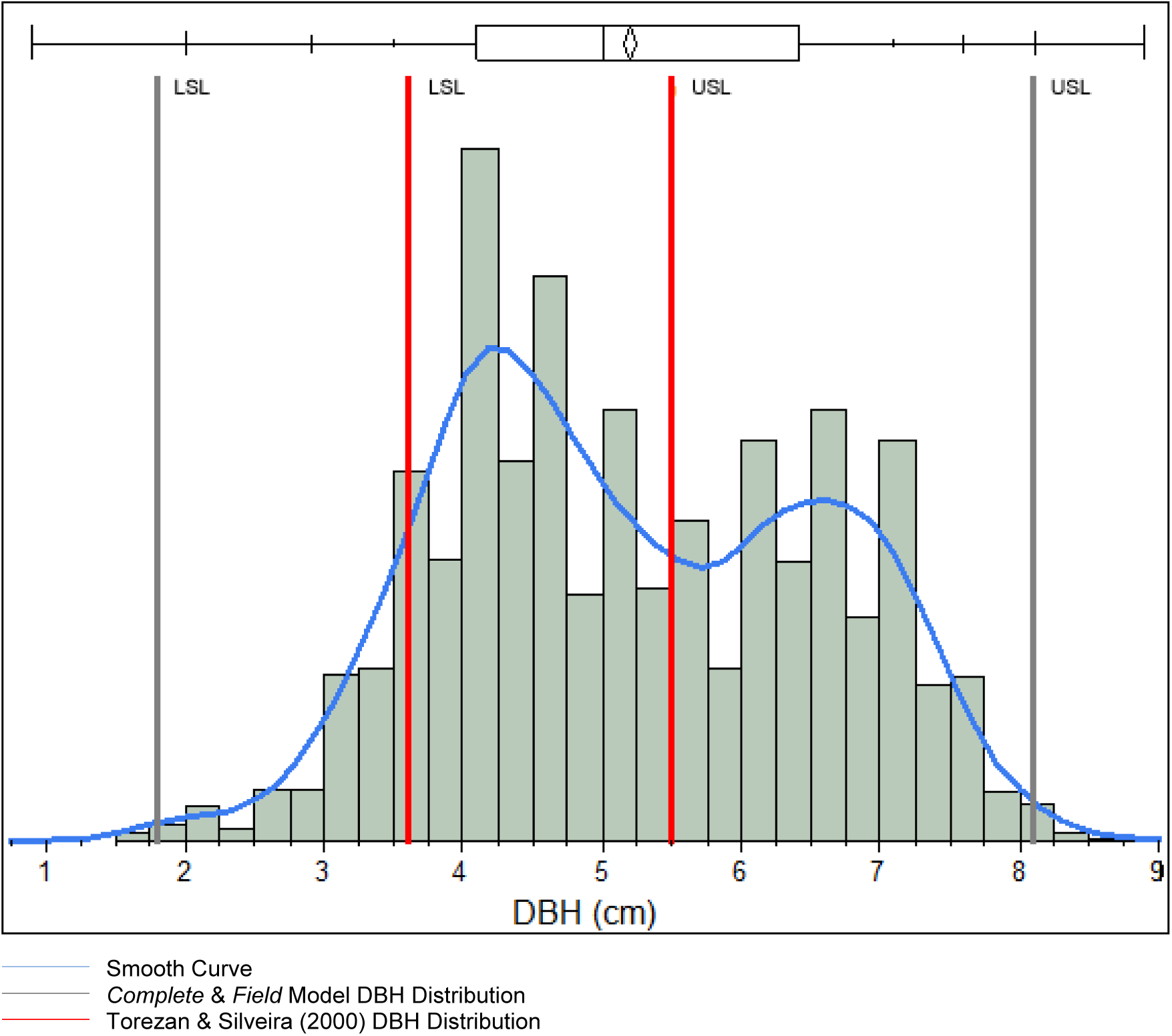
A non-parametric (kernel) distribution function fit to the 3,966 ramets measured within the ten 0.36-ha plots. Outer grey lower and upper specified limis (LSL and USL respectively) represent the range of size distributions incorporated in the derivation of the *complete* and *field* allometric models derived here. The interior LSL and USL represent the range of size distributions incorporated in the derivation of the previous allometric models created by Torezan and Silveira (2000).

**Appendex Formulae 1.** Linear (1a) and 3rd degree polynomial (1b) allometric equations as derived by Torezan and Silveira (2000) for mature *G. weberbaueri* (n = 10, DBH range from 4.2-5.5 cm)

*(1a) Estimated Biomass = (5.4922*DBH) – 19.516.*

*(1b) Estimated Biomass = (2.928*DBH*^*3*^*) – (37.554*DBH*^*2*^*) + (161.23* DBH) - 226.54*

**Appendex Formula 2.** DBH only model derived from the ramets sampled in the analysis here. P < 0.0001 for DBH, Adj. R^2^ = 0.79, RMSE = 1.90, n = 278.

*(2) Complete Dry Weight = −7.208 + 2.875*DBH*

## Tables

**Table 1.** *Complete*, *field* and DBH-only model parameter estimates calculated for the ≤ 5 model predictors included in the stepwise linear regression analyses

|  |  |  |  |  |  | Variable input range and significance |  |  |  |
| --- | --- | --- | --- | --- | --- | --- | --- | --- | --- |
| Name | Aboveground biomass model | n | Adj. $R^2$ | Overall $p$ Value | RMSE | X1: DBH (cm) | X2: $\geq 2$ cm branch (#) | X3: Length (m) | X4: Small branch (#) |
| *Complete Model* | $-6.00 + (1.72x_1) + (1.34x_2) + (0.25x_3) + (0.08x_4)$ | 192 | 0.93 | < 0.0001 | 1.24 | 1.8 - 8.1 *** | 0 - 7 *** | 0.34 - 24.81 *** | 0-32 *** |
| *Field Model* | $-6.24 + (2.54x_1) + (1.07x_2)$ | 278 | 0.84 | < 0.0001 | 1.66 | 1.8 - 8.1 *** | 0 - 7 *** | --- | --- |
| *DBH Model* | $-7.208 + 2.875x_1$ | 278 | 0.79 | < 0.0001 | 3.86 | 1.8 - 8.1 *** | --- | --- | --- |

\*, \*\*, \*\*\* indicate p values of 0.01, 0.001 and > 0.0001

## References

Asner GP, Powell GVN, Mascaro J, Knapp DE, Clark JK, Jacobson J, Kennedy--Bowdoin T, Balaji A, Paez-Acosta G, Victoria E, Secada L, Valqui M, Hughes RF. 2010. High-resolution forest carbon stocks and emissions in the Amazon. Proceedings of the National Academy of Sciences of the United States of America 107: 16738–16742.

Carvalho AM. 2009. Ciclo de vida de populações de bambu (Guadua spp.) no tempo e no espaço, no Sudoeste da Amazônia. Programa de Pós-Graduação em Ciências de Floresta Tropicais - CFT. Universidade do Amazonas, Instituto Nacional de Pesquisas da Amazonia-INPA, Manaus-AM, p. 60.

Carvalho AM, Nelson BW, Bianchini MC, Plagnol D, Kuplich TM, Daly DC. 2013. Bamboo-dominated forests of the southwest Amazon: detection, spatial extent, life cycle length and flowering waves. PLoS ONE 8.

Chave J, Andalo C, Brown S, Cairns MA, Chambers JQ, Eamus D, Folster H, Fromard F, Higuchi N, Kira T, Lescure JP, Nelson BW, Ogawa H, Puig H, Riera B, Yamakura T. 2005. Tree allometry and improved estimation of carbon stocks and balance in tropical forests. Oecologia 145: 87–99.

Conover A. 1994. A new world comes to life, discovered in a stalk of bamboo. Smithsonian 25: 120.

Cruz RH. 2009. Bambú - Guadua: Guadua angustifolia Kunth. Guadua angustifolia Kunth. Bosques Naturales en Colombia y Plantaciones Comerciales en México, Pereira, Risaralda, Colombia, p. 720.

Davidson EA, de Araujo AC, Artaxo P, Balch JK, Brown IF, Bustamante MMC, Coe MT, DeFries RS, Keller M, Longo M, Munger JW, Schroeder W, Soares BS, Souza CM, Wofsy SC. 2012. The Amazon Basin in transition. Nature 481: 321–328.

Doust LL. 1981. Population dynamics and local specialization in a clonal perennial (*Ranunculus repens)*: I. The dynamics of ramets in contrasting habitats. The Journal of Ecology69: 13.

França MB. 2002. Modelagem de biomassa florestal através do padrão espectral no Sudoeste da Amazônia. Programa de Pós-graduação em Biologia Tropical e Recursos Naturais. Universidade do Amazonas - UA, Instituto Nacional de Pesquisas da Amazônia - INPA, Manaus- AM, p. 106.

Griscom BW, Ashton PMS. 2003. Bamboo control of forest succession: *Guadua sarcocarpa* in Southeastern Peru. Forest Ecology and Management175: 445–454.

Griscom BW, Ashton PMS. 2006. A self-perpetuating bamboo disturbance cycle in a neotropical forest. Journal of Tropical Ecology22: 587–597.

Josse C, Navarro G, Encarnación F, Tovar A, Comer P, Ferreira W, Rodríguez F, Saito J, Sanjurjo J, Dyson J, Rubin de Celis E, Zárate R, Chang J, Ahuite M, Vargas V, Paredes F, Castro W, Maco J, Arreátegui F. 2007. Digital Ecological Systems Map of the Amazon Basin of Peru and Bolivia. In: NatureServe (Ed.), Arlington, VA, USA.

Judziewicz EJ, Clark LG, Londono X, Stern MJ. 1999. American bamboos. Smithsonian Institution Press, Washington D.C.; London, England.

Jyoti Nath A, Das G, Das AK. 2009. Above ground standing biomass and carbon storage in village bamboos in North East India 1188-1196. Biomass and Bioenergy 33.9: 9.

Kitani O, Hall CW. 1989. Biomass handbook. Gordon and Breach Science Publishers, New York, NY.

Kumar BM, Rajesh G, Sudheesh KG. 2005. Aboveground biomass production and nutrient uptake of thorny bamboo [Bambusa bambos (L.) Voss] in the homegardens of Thrissur, Kerala. Journal of Tropical Agriculture43: 6.

Londoño X, Peterson PM. 1991. Guadua sarcocarpa (Poaceae, Bambuseae), a new species of Amazonian bamboo with fleshy fruits. Systematic Botany 16: 630–638.

Louton J, Gelhaus J, Bouchard R. 1996. The aquatic macrofauna of water-filled bamboo (Poaceae: Bambusoideae: Guadua) internodes in a Peruvian lowland tropical forest. Biotropica 28: 228–242.

McClure FA. 1973. Bambusa weberbaueri (Pilger). In: Soderstrom TR (Ed.), Smithsonian Contribution to Botany. Smithsonian Institution, Washington D.C., p. 148.

Nelson BW, Alves de Oliveira AC, Batista GT, Vidalenc D, Silveira M. 2001. Modeling biomass of forests in the southwest Amazon by polar ordination of Landsat TM. In: INPE (Ed.), Anais XSBSR, Foz do Iguacu, Brasil, pp. 1683–1690.

Nelson BW, de Oliveira ACA, Vidalenc D, Smith M, Bianchini MC, Nogueira EM. 2006. Florestas dominadas por bambus semi-escandentes do gênero Guadua, no sudoeste da Amazônia. In: Brasília Ud (Ed.), Anais do Seminário Nacional de Bambu, Brasilia D.F., Brasil, pp. 49–55.

Nelson BW, Kalliola R, Shepard G. 1997. Tabocais de Guadua spp. no sudeste Amazônico: extensão geográfica, mortalidade sincronizada e relação com incêndios florestais. 38th National Botanical Congress. Sociedade Botânica do Brasil, Universidade Regional do Cariri; Crato-Ceara, p. 163.

Niklas KJ. 2004. Plant allometry: is there a grand unifying theory? Biol Rev Camb Philos Soc 79: 871–889.

Nogueira EM, Fearnside PM, Nelson BW, França MB. 2007. Wood density in forests of Brazil’s ’arc of deforestation’: Implications for biomass and flux of carbon from land-use change in Amazonia. Forest Ecology and Management248: 119–135.

Nogueira EM, Nelson BW, Fearnside PM, França MB, Oliveira ÁCAd. 2008. Tree height in Brazil’sarc of deforestation’: Shorter trees in south and southwest Amazonia imply lower biomass. Forest Ecology and Management255: 2963–2972.

Oliveira ACA. 2000. Efeitos do bambu *Guadua weberbaueri* Pilger sobre a fisionomia e estrutura de uma floresta no Sudoeste da Amazonia. Programa de Pós Graduação em Biologia Tropical e Recursos Naturais. Universidade do Amazonas – UA, Instituto Nacional de Pesquisas da Amazonia – INPA, Manaus-AM, p. 71.

Olivier J. 2008. Gramíneas (Poaceae) bambusiformes del Río de Los Amigos, Madre de Dios, Perú. Revista Peruana de Biología 15: 6.

Olivier J, Otto T, Roddaz M, Antoine PO, Londono X, Clark LG. 2009. First macrofossil evidence of a pre-Holocene thorny bamboo cf. Guadua (Poaceae: Bambusoideae: Bambuseae: Guaduinae) in south-western Amazonia (Madre de Dios - Peru). Review of Palaeobotany and Palynology153: 1–7.

Osorio JA, Velez JM, Ciro HJ. 2007. Internal structure of the Guadua and its incidence in the mechanical properties. Dyna-Colombia 74: 81–94.

Regard V, Lagnous R, Espurt N, Darrozes J, Baby P, Roddaz M, Calderon Y, Hermoza W. 2009. Geomorphic evidence for recent uplift of the Fitzcarrald Arch (Peru): A response to the Nazca Ridge subduction. Geomorphology 107: 107–117.

Riano NM, Londono X, Lopez Y, Gomez JH. 2002. Plant growth and biomass distribution on Guadua angustifolia Kunth in relation to ageing in the Valle del Cauca - Colombia. Bamboo Science and Culture: The Journal of the American Bamboo Society 16: 9.

Saatchi SS, Houghton RA, Alvala R, Soares JV, Yu Y. 2007. Distribution of aboveground live biomass in the Amazon basin. Global Change Biology 13: 816–837.

Salimon CI, Putz FE, Menezes L, Anderson A, Silveira M, Brown IF, Oliveira LC. 2011. Estimating state-wide biomass carbon stocks for a REDD plan in Acre, Brazil. Forest Ecology and Management262: 555–560.

Silman MR, Ancaya EJ, Brinson J. 2003. Bamboo forests of western Amazonia. In: Leite R, Pitman N, Alvarez P (Eds.), Alto Purus: Biodiversity, Conservation and Management. Duke University Center for Tropical Conservation Press, Durham, NC, pp. 63–72.

Silveira M. 1999. Ecological aspects of bamboo-dominated forest in southwestern Amazonia: An ethnoscience perspective. Ecotropica 5: 4.

Smith M. 2000. Efeito de perturbações sobre a abundância, biomassa e arquitetura de Guadua weberbaueri Pilg. (Poaceae – Bambusoideae) em uma floresta dominada por bambu no Sudoeste da Amazônia. Programa de Pós-Graduação em Biologia Tropical e Recursos Naturais. Universidade do Amzonas (UA), Instituto Nacional de Pesquisas da Amazonia (INPA), Manaus-AM, p. 80.

Smith M, Nelson BW. 2011. Fire favours expansion of bamboo-dominated forests in the south-west Amazon. Journal of Tropical Ecology27: 59–64.

Torezan JMD, Silveira M. 2000. The biomass of bamboo (Guadua weberbaueri Pilger) in open forest of the southwestern Amazon. Ecotropica (Bonn) 6: 71–76.

UNFAO. 2005a. Global forest resources assessment-Brazil country report on bamboo resources. In: Rattan INfBa (Ed.).

UNFAO. 2005b. Global forest resources assessment: progress towards sustainable forest management. Food and Agriculture Organization of the United Nations, Rome.

v-c-s.org. 2011. The verified carbon standard project database. In: Association V (Ed.).

Vidalenc D. 2000. Distribuição das florestas sominadas pelo bambu Guadua weberbaueri em escala de paisagem no sudoeste da Amazônia e fatores edáficos que afetam sua densidade. Universidade do Amazonas-UA, Instituto Nacional de Pesquisas da Amazonia-INPA, Manaus-AM, p. 95.

Zhou GM, Meng CF, Jiang PK, Xu QF. 2011. Review of carbon fixation in bamboo forests in China. Botanical Review 77: 262–270.

